# Cortical organoids reveal human-specific roles of METTL5 in neurodevelopment via regulation of CHCHD2

**DOI:** 10.1101/2025.07.13.664555

**Authors:** Elena M Turkalj, Mariami Kuljanishvili, Gugene Kang, Isabelle Liu, Chloe Ghent, Carlo Waltier Soriano, Neda Keyhanvar, Daniel Brody, Michael Oldham, Caroline Vissers

**Affiliations:** University of California, San Francisco

## Abstract

METTL5 is a conserved methyltransferase responsible for catalyzing N6-methyladenosine (m^6^A) modification on 18S ribosomal RNA. Human patients with mutations in *METTL5* show severe microcephaly and intellectual disability, which has not been fully recapitulated in animal models. Given its emerging role in neurodevelopment, we sought to investigate METTL5 function in a human-specific context using cortical forebrain organoids derived from human induced pluripotent stem cells. We generated *METTL5* knockout organoids and observed a marked delay in neural stem cell proliferation and the timing of neuronal differentiation, suggesting a critical role for METTL5 in the temporal regulation of neurogenesis. Though METTL5 methylates rRNA near the decoding center, the mechanism of this methylation remains highly contested. In our cortical organoids, broad translational changes mirror stress response rather than transcript-specific regulation of translation. Transcriptomic analysis further revealed a significant downregulation of *CHCHD2*, a nuclear-encoded mitochondrial gene linked to cellular energy metabolism and neurodevelopmental processes. Overexpression of CHCHD2 rescued proliferation defects of *METTL5*-KO neural progenitor cells, suggesting dysregulation of CHCHD2 is heavily involved in the phenotypes of *METTL5* mutant patient brains. These findings highlight a previously uncharacterized link between METTL5, ribosomal RNA modification, and cellular metabolism essential for proper human brain development.

## Introduction

The epitranscriptome is rapidly gaining interest as a crucial regulator of gene expression regulation across development and disease. Hundreds of different modifications occur across mRNA, tRNA, and rRNA, and disentangling the methyltransferases and individual functions of each modification has been challenging. In 2019, METTL5 was identified as the enzyme responsible for installing N6-methyladenosine (m^6^A) on a single position (A1832) in 18S ribosomal RNA. The m^6^A_1832_ modification occurs just a few nucleotides away from the ribosome decoding center, suggesting a role in regulation of translation (*1, 2*).

Human genetics studies of METTL5 have identified two bi-allelic frameshift mutations (NM_014168.2:c.344_345delGA, NM_014168.2:c.571_572delAA) (*3*), a homozygous SNP in the methyltransferase domain of exon 3 (NM_014268.3:c362A>G) (*4*), a homozygous deletion mutation in the donor splice site of exon 2 (NM_014168:c.223_224+8del) (*5*), and a homozygous intron mutation (NM_014168.4: c.224+5 G > A) (*6*) that causes exon 2 skipping . All of these mutations cause intellectual disability (ID) and microcephaly, with variable additional symptoms including seizures, craniofacial defects, growth retardation, and aggression. The average severity of microcephaly from these studies is 3.6 standard deviations below the mean for the respective gestational ages, meaning it is categorized as severe microcephaly (*7*). Surprisingly, of the four studies published using Mettl5 knockout (KO) mouse models, microcephaly is not a clear or consistent phenotype (*8–11*). Micro-CT images of mouse skulls at 6 and 14 weeks do not show clear reductions in brain size, though circumference and volume are not explicitly measured (*8, 9*). Instead, growth defects in body weight and craniofacial abnormalities are pervasive in Mettl5-KO mice, and behavioral tests for intellectual disability show significance in (*11*) and (*8*), but not in (*10*). Finally, *METTL5-KO* in zebrafish did show a microcephaly phenotype, with brain size reductions occurring in the forebrain and midbrain, but not hindbrain, of knockout animals (*3*). Overall, the inconsistency in phenotype in mice and across species suggests there may be cell-type or species specificity of METTL5 function. In this study we generate *METTL5* knockout human induced pluripotent stem cells (hiPSCs) and use these to model human neural development with hiPSC-derived cortical forebrain organoids. This is the first human-specific, 3D model of *METTL5* function, and specifically highlights the importance of *METTL5* in human neural development.

The mechanisms of action of METTL5 have been largely focused on regulation of translation due to the precarious positioning of m^6^A_1832_ near the ribosomal decoding center. However, work over the past five years has shown that loss of METTL5 does not affect ribosomal biogenesis (*3, 10*) or rRNA maturation (*12*). Reports on whether changes in translation are global or transcript-specific have largely conflicted. Global reductions in translation upon *METTL5* loss were reported in HEK293 and mESCs, while transcript-specific changes were reported in mESCs, Hela, HepG2, and Hct116 cells (*8, 10, 13–15*). This conflicting data again suggests there may be certain contexts in which *METTL5* is especially important, further pushing the need for studies that directly model the most severe phenotypes seen in humans.

Here, we show that *METTL5* is highly expressed in radial glial cells along the ventricular structures of hiPSC-derived forebrain organoids, and neural progenitor cells (NPCs) are profoundly impaired in proliferation and neural differentiation upon knockout of *METTL5*. We find a reduction in translation with no clear evidence for transcript specificity. Instead, single cell RNA sequencing (sc RNA-seq) reveals highly significant downregulation of *CHCHD2*, a gene involved in mitochondrial function and cellular respiration. To this end, we show that *METTL5-KO* NPCs cannot properly recover from mitochondrial stress, and overexpression of *CHCHD2* rescues the proliferation defects of *METTL5-KO* NPCs. Recent work has poised *CHCHD2* as an emerging key gene involved in multiple types of human neurodevelopmental and neurodegenerative disorders (*16–20*). Some of the severity of brain phenotypes seen in *METTL5* mutant human patients could be due to downstream effects of *CHCHD2* dysregulation during neural development. Altogether, we propose that *METTL5* is especially important in contexts where cells have a high metabolic demand–such as in the developing human brain– and that indirect downstream effects on key genes such as *CHCHD2* may drive the specific neural phenotypes and severities seen in humans.

## Results

We generated *METTL5* knockout hiPSCs/ESC for three different background lines: KOLF2.1j, H9, and WTC11. We used three unique gRNAs in concert targeting exon 1 of *METTL5* (**Figure 1A**), and isolated multiple clones for each cell line. We performed qPCR validation of *METTL5* knockout using primers on exon 1 (**Figure 1B**), then performed a Western blot to confirm loss of METTL5 protein (**Figure 1C**). Next, we confirmed loss of m^6^A at the A1832 site in 18S rRNA by purifying a 40-nucleotide fragment around the A1832 site and performing an m^6^A ELISA (**Figure 1D**).

**Figure 1:**
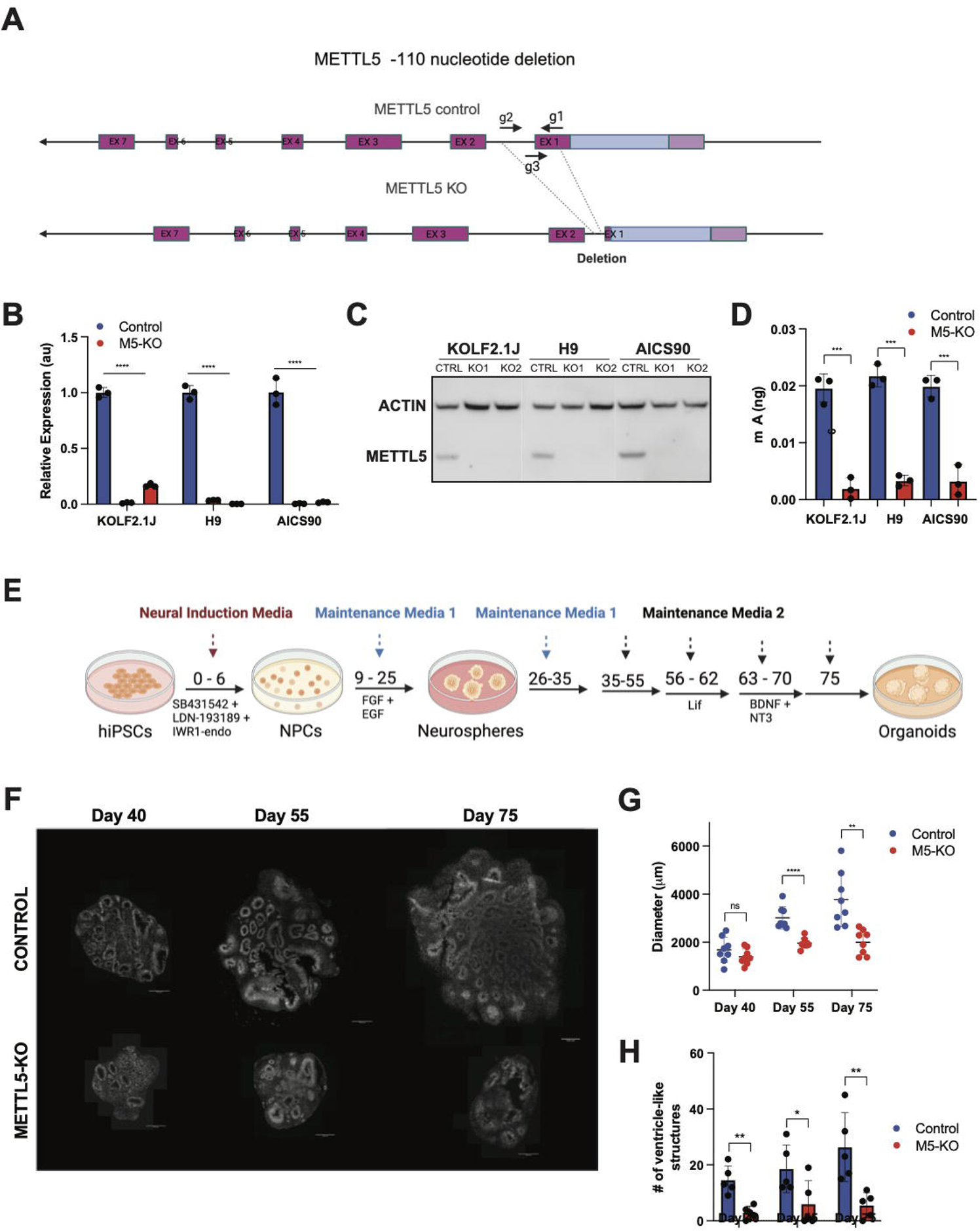
METTL5 Knockout hiPSCs lack m^6^A_1832_ and recapitulate microcephaly in Day 40, 55, and 75 forebrain cortical organoids. **A.** gRNAs designed for METTL5 knockout targeting the first exon with a 110 bp deletion that causes a frameshift and disrupts the ORF. **B.** qPCR validation of *METTL5* KO in two distinct KO clones for each KOLF2.1J, H9, and AICS90 pluripotent stem cell lines shows loss of *METTL5* transcript in KOs; **** p<0.0001. **C.** Western blot confirmation of METTL5 protein loss in all KO cell lines. **D.** m^6^A ELISA shows significant reduction in m^6^A from twice purified 40-nt fragments of 18S rRNA containing A_1832_. n=3 replicates; ***p<0.001. **E.** Cortical forebrain organoid differentiation protocol starting from hiPSCs and growing for 75 days. **F-H.** DAPI-stained KOLF2.1J Control and *METTL5-*KO organoids at Days 40, 55, and 75 show significant differences in organoid diameter and the number of ventricle-like structures per organoid. n=5-8 organoids/timepoint for control and KO; ns= no significance, *p<0.05, ** p<0.01, ***p<0.001, ****p<0.0001.

In order to model human neural development, we grew cortical forebrain organoids using an adapted protocol from (*21–24*) (**Figure 1E**). Briefly, we used dual SMAD inhibition for days 0-6 to drive neural progenitor formation in a Slitwell plate (*24*). Then we added EGF and FGF for days 9-25 to generate neurospheres; we moved the organoids to a low attachment 6 well plate on an orbital shaker at day 18. We then cultured the organoids in maintenance media without Vitamin A from days 26-35, then switched to maintenance media with Vitamin A for the remainder of the protocol. Lif was added at days 56-62, and BDNF and NT3 were added at days 63-70. Organoids were collected at days 40, 55, and 75. *METTL5* knockout caused significant reductions in the number of ventricle-like structures at Days 40, 55, and 75, and significant reductions in diameter at Days 55 and 75. (**Figure 1F-H**). H9 and AICS90-derived organoids confirmed these findings at Day 40 (**Supplementary Figure 1**). KOLF2.1J organoids were more neurogenic in terms of ventricle-like structure formation compared to both H9 and AICS90. Therefore, we continued our organoid studies focusing on the KOLF2.1J hiPSC line.

### METTL5 is highly expressed in radial glial cells (RGCs) and drives the differentiation schedule of cortical development

Immunohistochemical staining show strong METTL5 staining in NESTIN-positive RGCs along the inner walls of the ventricular-like structures of our organoid models (**Figure 2A**). This particular staining highlights the utility of human brain organoid models, since sub-ventricular RGCs are not easily grown in 2D. Interestingly, Mettl5 was not seen in embryonic mouse day 17.5 Nestin+ sub-ventricular RGCs (**Supplementary Figure 2**), suggesting that METTL5 may have human-specific impacts on neural development.

**Figure 2:**
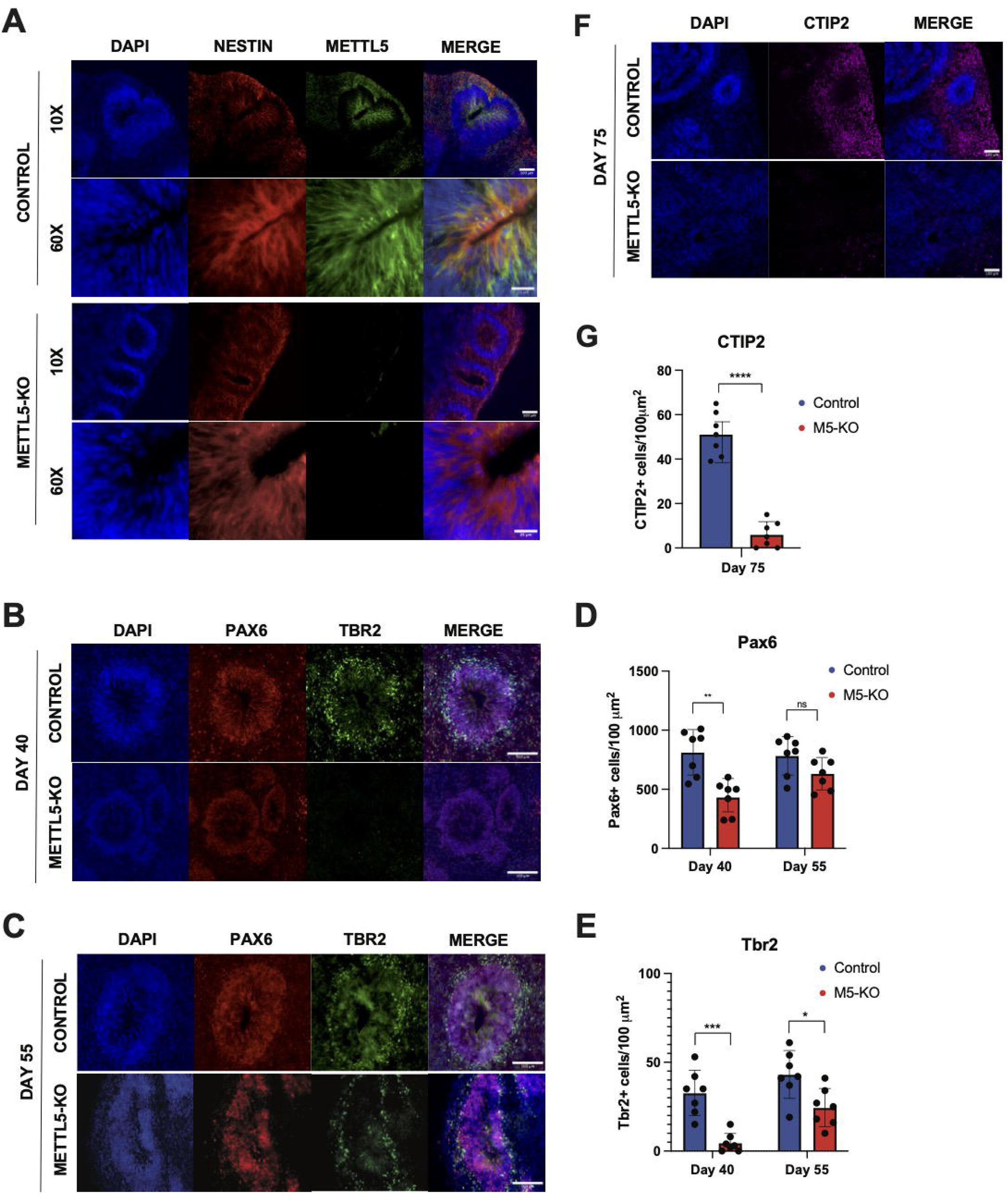
METTL5 is highly expressed in ventricular zone radial glial cells and METTL5-KO organoids show significant developmental delays. **A.** Day 55 organoids show strong METTL5 expression in Nestin+ radial glial cells (top), which is lost in METTL5-KO organoids (bottom). **B.** Day 40 control organoids show thick Pax6+ zones around ventricle-like neural rosettes with Tbr2+ intermediate progenitor cells (IPCs) emerging at the edges. METTL5-KO organoids have reduced numbers of Pax6+ cells and have not yet started producing Tbr2+ IPCs. Scale bar = 100µm **C.** Day 55 control and METTL5-KO organoids show increased Tbr2+ IPC production in the control relative to Day 40, equal levels of Pax6+ cells between KO and control, but fewer Tbr2+ cells in KO versus control. Scale bar = 100 µm. **D,E.** Quantification of Pax6+ and Tbr2+ cells per 100 µm^2^ area tangential to ventricular wall at Day 40 and 55. n=7. ns= no significance. *p<0.05, ** p<0.01, ***p<0.001. **F.** Day 75 control organoids show robust CTIP2+ neuron production, while METTL5-KO organoids have significantly fewer, quantified in **G.** n=7, ****p<0.0001.

### Loss of *METTL5* causes impairments in early neurogenesis and the formation of subventricular-like zones

Ventricular-like structures are defined by a thick layer of Pax6+ neural progenitor cells (NPCs) around a ventricle, and an outer layer of Tbr2+ intermediate progenitor cells (IPCs). At Day 40, *METTL5* knockout organoids show significantly fewer Pax6+ NPCs per 100µm^2^ of ventricular area relative to controls, and have not yet started producing Tbr2+ IPCs (**Figure 2B**). By Day 55, knockout organoids catch up with controls in the number of Pax6+ cells and have started producing Tbr2+ IPCs. However, KOs still have significantly fewer Tbr2+ IPCs compared to controls at Day 55 (**Figure 2C-E**). This delay is continued into neurogenesis; at Day 75, control organoids show significant CTIP2+ neuron populations, while knockout organoids show significantly reduced numbers of CTIP2+ neurons (**Figure 2F-G**). These data suggest a developmental delay upon loss of *METTL5*, without a complete loss of stem cell competence to differentiate down the neural lineage.

### *METTL5* drives RGC proliferation and differentiation in 3D and 2D

The significant reduction in *METTL5-*KO organoid diameter and the reduction in Pax6+ cells at Day 40 suggests a deficit in proliferation capacity of *METTL5*-KO NPCs. Indeed, Day 55 *METTL5-*KO organoids show significant reductions in PH3+ cells undergoing mitosis along the walls of the ventricle-like structures and actively cycling Ki67+ cells (**Figure 3A-B**). Though cell counts were normalized to 100µm^2^ of ventricular area, the difference in size of ventricle-like structures may account for some differences in the number of proliferating cells. So, we differentiated control and *Mettl5-*KO hiPSCs into NPCs (*25, 26*) and quantified NPC proliferation rate in 2D. *METTL5-*KO NPCs showed significantly reduced proliferation rates relative to controls in all cell lines tested (**Figure 3C, Supplementary Figure 3A-B**).

**Figure 3:**
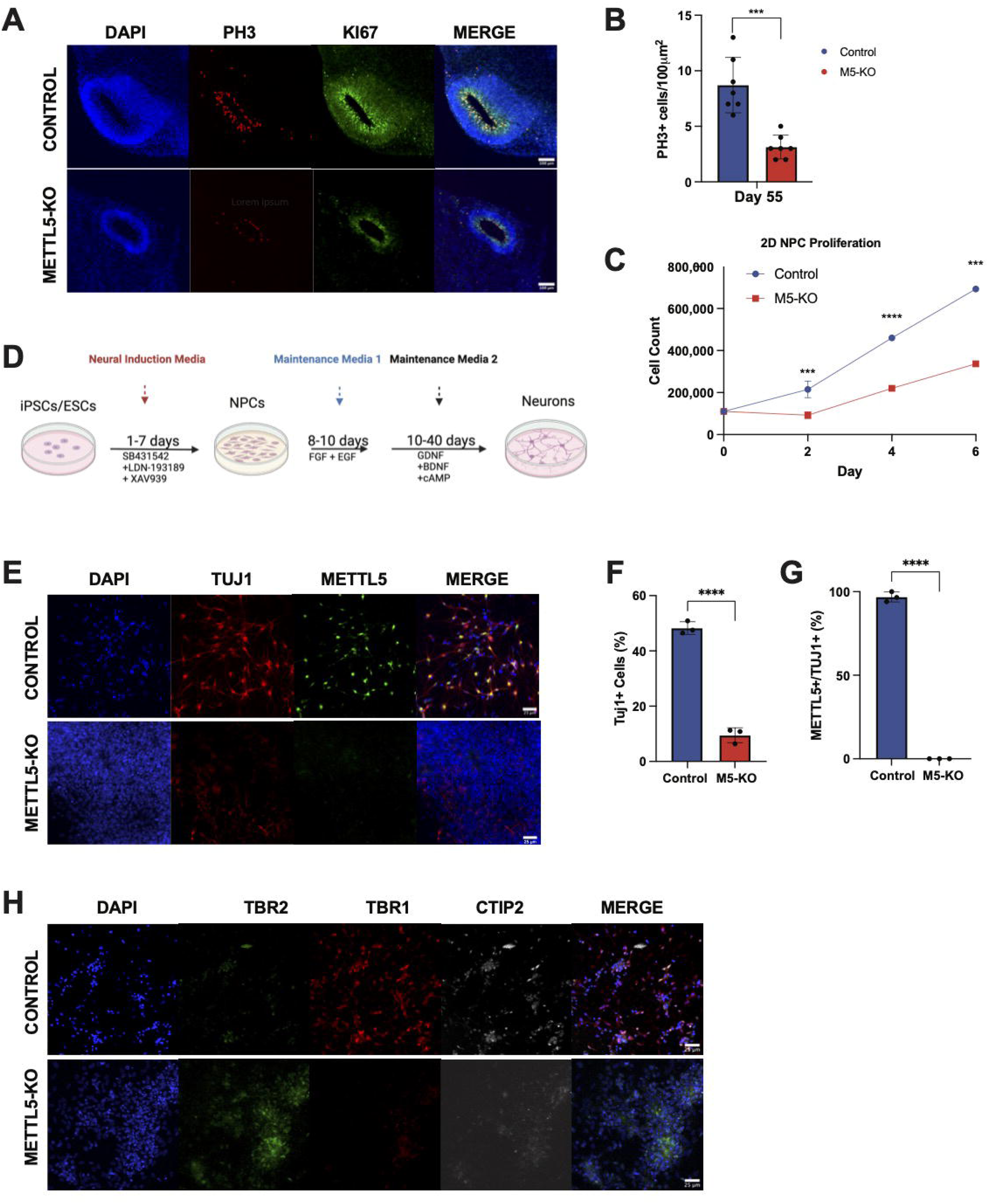
Proliferation and neural differentiation are significantly impaired in METTL5-knockout neural progenitor cells (NPCs). **A.** Day 55 organoids show a significant reduction in PH3+ cells and Ki67+ cells around ventricle-like structures. Scale bar=100µm **B.** Quantification of PH3+ cells per 100 µm^2^ area tangential to ventricular wall in control versus METTL5-KO organoids. n=7 ventricular structures. ***p<0.001. **C.** Control and METTL5-KO hiPSC were differentiated into NPCs, which were tested for proliferation rate by counting cells over 6 days after seeding equal numbers. Control NPCs show significantly faster proliferation than METTL5-KO NPCs, measured 2, 4, and 6 days after seeding. n=3 wells per timepoint; ***p<0.001, ****p<0.0001. **D.** 2D neural differentiation protocol to generate neurons from hiPSCs. **E.** Day 20 of neural differentiation shows significantly more TUJ1+ neurons, which are largely also METTL5+, in control cells relative to METTL5-KO cells. Scale bar = 25µm. **F-G.** Quantification of TUJ1+ cells and METTL5+/TUJ1+ cells from (**E**); n=3, ****p<0.0001. **H.** At Day 30 of neural differentiation control cells show high levels of Tbr1+ and CTIP2+ neurons and low levels of Tbr2+ IPCs. In contrast, METTL5-KO cells still express Tbr2+ with minimal Tbr1+ or CTIP2+ cells. Scale bar = 25µm.

Next, we performed 2D neural induction (*25, 26*) from hiPSCs to form TUJ1+ neurons (**Figure 3D**). By Day 20, control cells had formed neurons and were highly positive for both METTL5 and TUJ1. In comparison, *METTL5-*KO hiPSCs showed significantly reduced TUJ1+ staining and maintained NPC-like morphology and density in culture (**Figure 3E-G**). By Day 30, control cells were positive for deep layer markers CTIP2 and TBR1, while *METTL5-KO* cells were negative for neuronal markers, but positive for the earlier intermediate progenitor cell marker Tbr2 (**Figure 3H**). This largely recapitulates the delay in neurogenesis seen in our organoid models. To ensure that the reductions in proliferation and differentiation upon loss of *METTL5* were not due to increases in apoptosis, we performed a cleaved Caspase 3 assay. There were no significant differences in apoptosis between control and *METTL5-KO* NPCs at baseline, though addition of staurosporine as a positive control did elicit a larger apoptotic response in *METTL5*-KO NPCs compared to controls (**Supplementary Figure 3C-D**).

### METTL5 regulates global translation levels in NPCs and iPSCs

Previous studies on the mechanism of *METTL5* function reached conflicting conclusions on whether it regulates global or transcript-specific changes in translation. We found no indication that any particular subset of transcripts was singularly driving the phenotypes seen in NPCs and brain organoids. To quantify changes in global levels of translation, we performed polysome profiling on NPCs and iPSCs. There is a large increase in the monosome peak of METTL5-KO NPCs, which translates into a 57% decrease in the polysome-to-monosome (P/M) AUC ratio relative to control cells (**Figure 4A**). Although the absolute polysome peaks appear similar between KO and control NPCs, this is typical for neural progenitor profiles because total translation often declines as cells differentiate away from the highly translational state of iPSCs. Indeed, global protein synthesis rises briefly when pluripotent cells exit self-renewal and then falls as they adopt a neural progenitor or neuronal identity (*27*). To confirm that the monosome build-up reflects a genuine loss of translating ribosomes, we repeated the profiling in hiPSCs. In this setting, the METTL5-KO trace showed a pronounced loss of heavy polysomes, yielding an approximately 70 % reduction in polysome area (and a drop in the P/M ratio) relative to control cells (**Supplementary Figure 4A**).

**Figure 4:**
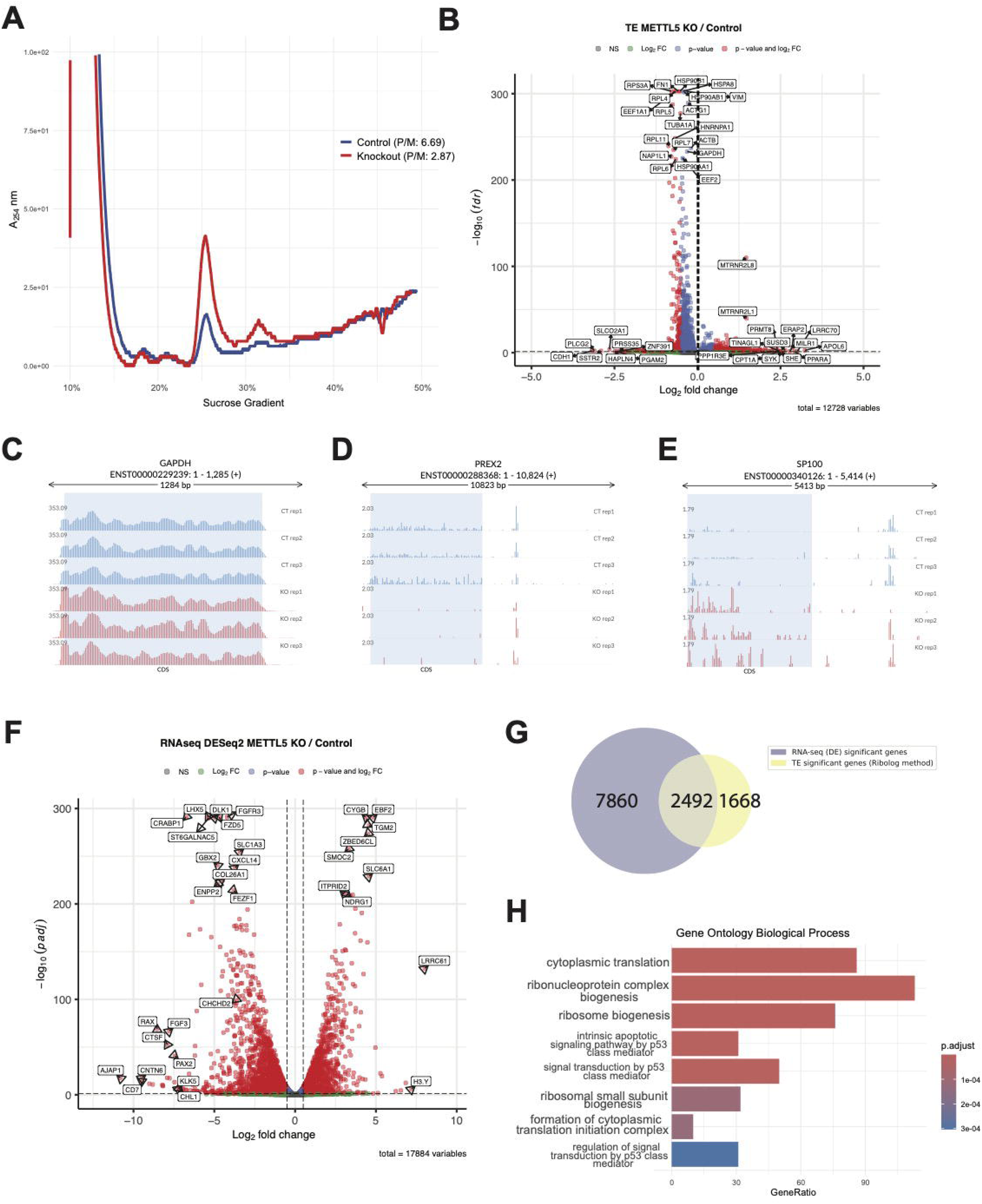
Translation is globally downregulated in METTL5-KO NPCs. **A.** Polysome profile shows a large increase in the monosome peak of METTL5-KO NPCs compared to control, though there was no clear change in polysomes, which were low in both control and knockout NPCs. Polysome/monosome ratios are 6.69 and 2.87 for control and knockout, respectively. **B**. Translation efficiency measured by Ribosome sequencing and processing with RiboLog shows many transcripts with downregulated translation efficiency in METTL5-KO NPCs relative to control. Transcripts with Log2 fold-change > 0.5 or < -0.5 and FDR<0.05 shown in red; remaining transcripts with FDR<0.05 but Log2FC below +/-0.5 shown in blue. Genes with Log2FC >0.5 or < -0.5 but FDR>0.05 shown in green. **C.** Sample coverage plot from Ribo-Seq for control gene GAPDH with no change in TE, **D.** PREX2 coverage plot shows significantly fewer ribosome protected fragments in the coding sequence in *METTL5-*KO NPCs across replicates, and **E.** SP100 coverage plot exemplifies transcripts with increased translation in *METTL5-*KO NPCs compared to controls. **F.** RNA sequencing done on the same samples used for Ribo-seq processed for differential expression analysis show thousands of up-and down-regulated transcripts in METTL5 KO NPCs versus control. **G.** Quantification of the number of differentially regulated transcripts at the transcriptional level (purple, 7860), the translational level (yellow, 1668), and both (gray, 2492). **H.** Gene ontology analysis for biological processes in the 2492 genes differentially regulated at both the transcriptional and translational levels show significant enrichment for multiple terms related to translation and ribosomal biogenesis.

To further probe the effects of METTL5 loss on translation efficiency, we performed Ribosome sequencing on control and *METTL5-KO* NPCs and used the Ribolog method to quantify translation efficiency (*28*). Overall, there was a downward shift in translation efficiency of the KO, with 2936 transcripts (23.1%) showing a significant reduction in translation efficiency and 1224 transcripts (9.6%) showing an increase in translation efficiency compared to controls (**Figure 4B-E, Supplementary Figure 4B**). Motif analysis showed enrichment for general GC-rich repeats in both the up-and down-regulated translated transcripts (**Supplementary Figure 4C-D**), suggesting there is no singular motif or subset of transcripts that is exclusively susceptible to changes in m^6^A on A_1832_. However, narrowing our analysis to the 100 most significantly downregulated transcripts from our Ribo-log dataset revealed a significant enrichment of a pyrimidine-rich PRTE motif, which was present in the 5’ UTRs of 21 out of the 100 transcripts. (**Supplementary Figure 4E**). PRTE motifs are known modulators of translation efficiency, often found in the 5’ UTRs of transcripts regulated by nutrient availability and cellular stress. They have been implicated in mTORC1-sensitive translational control by making transcripts more susceptible to repression under stress conditions. Therefore, some changes in translation efficiency seen in the knockout may be downstream effects of broader cellular stress, rather than direct interaction with m^6^A_1832_. Disentangling broad translational responses to cellular stress as opposed to direct effects of loss of m^6^A_1832_ in *METTL5-*KO cells remains difficult and is likely a cause of the conflicting conclusions of previous studies.

Our Ribo-seq analysis incorporated RNA-seq of input RNA from Control and *METTL5-*KO NPCs. Differential expression analysis on this RNA-seq data shows that 10,452 genes are significantly differentially expressed in KOs versus Control NPCs (**Figure 4F**). In comparison, only 4,160 gene transcripts have significant changes in translation efficiency, with 2,492 genes showing significant changes at both the transcriptional and translational levels (**Figure 4G**). These 2,492 genes are highly enriched for gene ontology categories involving translation and ribosomal biogenesis, suggesting a potential feedback loop that reinforces changes in translation at a global level (**Figure 4H**). There was no clear trend on correlation between directionality of transcriptional regulation and translation efficiency regulation (**Supplementary Figure 4F**).

### **Single cell RNA sequencing (scRNA-seq) shows differential organoid composition of** *METTL5-*KO and Control Day 75 organoids

Using scRNA-seq of Day 75 cortical organoids and labeling individual cell clusters by Seurat clustering and gene set enrichment analysis generates 16 unique cell clusters shared across WT and KO organoids (**Figure 5A, Supplementary Figure 5**). *METTL5*-KO organoids have proportionally more metabolically stressed cells and fewer RGCs (**Figure 5B**), which coincides with the reduction in organoid diameter and *METTL5*-KO NPC proliferation capacity seen in Figures 1-3.

**Figure 5:**
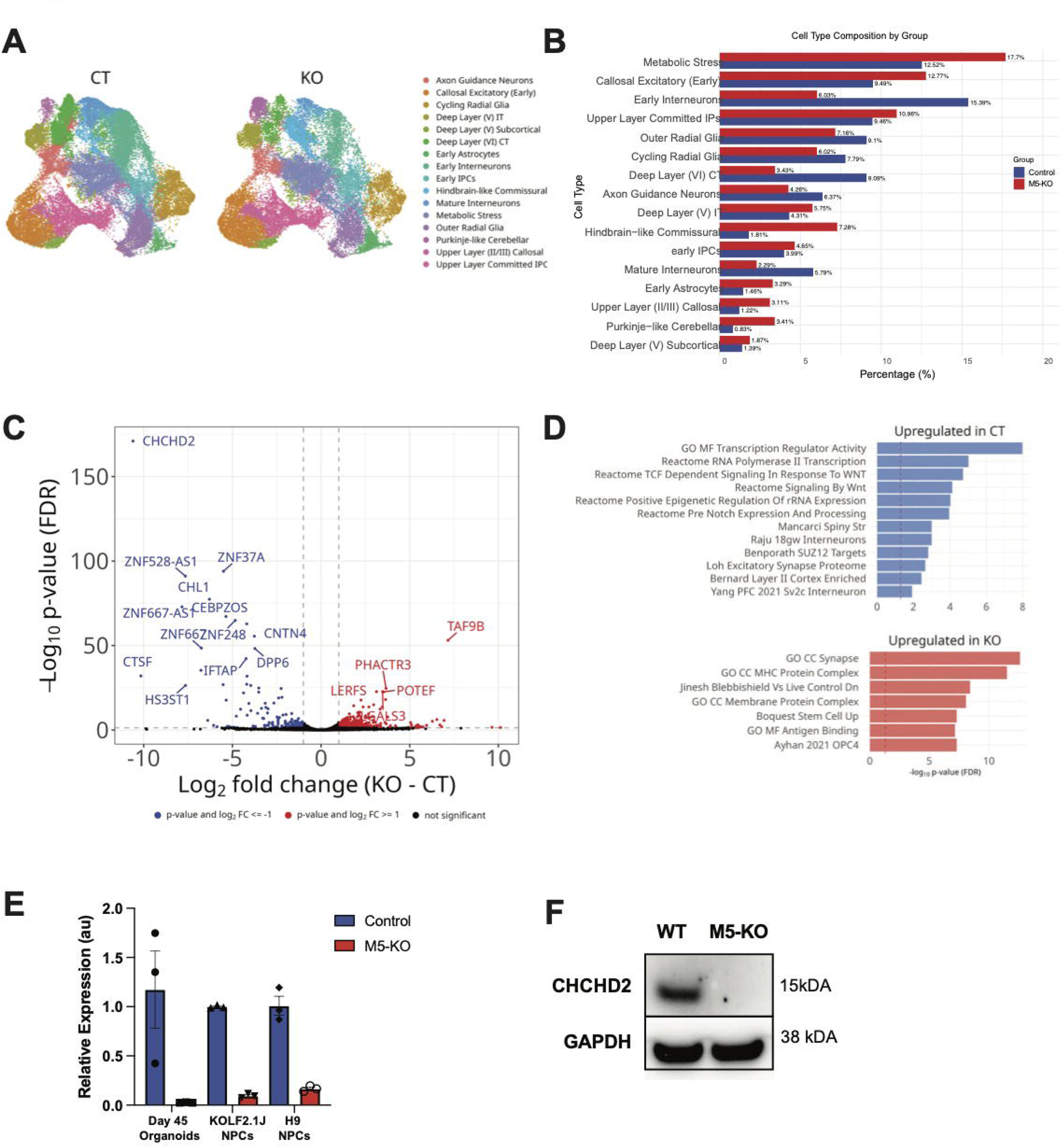
scRNA sequencing of Day 75 organoids highlight substantial changes in transcription in *METTL5-*KO organoids, particularly downregulation of neurogenic and mitochondrial regulatory gene CHCHD2. **A**. UMAPs of control (CT) and knockout (KO) nuclei, labeled by Seurat cluster. **B**. Proportions of nuclei from CT or KO for each cluster shown in (A). **C**. Volcano plot of differentially expressed genes between CT and KO. **D**. Enrichment analysis (one-sided Fisher’s exact test) of genes from (C) that are significant and have log_2_ fold change > 1 using published cell-type markers. P-values are FDR-corrected; dotted line represents significance at p < 0.05.

Using pseudo-bulk to perform differential expression analysis, we identified 4,532 genes that are differentially expressed in *METTL5-*KO Day 75 organoids (**Figure 5C**). Gene set enrichment and gene ontology analysis highlight regulation of transcription and Wnt signaling as upregulated in the Control versus *METTL5-*KO organoids, and Synapse and MHC Protein Complex cellular components as most upregulated in *METTL5-*KO organoids versus Controls (**Figure 5D**).

### CHCHD2 is massively downregulated in *METTL5-*KO Organoids and NPCs

The differential expression analysis both in pseudobulk and individual cell type clusters shows highly significant downregulation of CHCHD2 in *METTL5-*KO cells (**Supplementary Figure 6 & 7**). CHCHD2 is a gene that regulates mitochondrial function and the mitochondrial integrated stress response, with known involvement in neurodevelopmental processes (*29, 30*). We validated CHCHD2 downregulation via qPCR (**Figure 5E**) and Western blot (**Figure 5F**). Notably, CHCHD2 was not identified as a differentially expressed gene in *Mettl5*-KO mouse postnatal brains (*10, 11*). In parallel, CHCHD2 has recently been identified as involved in human-specific neurodevelopmental disorders such as Autism (*17*), and is significantly downregulated in neurons derived from hiPSCs of patients with lissencephaly (*16*).

### *METTL5-*KO cells are unable to respond to mitochondrial stress

Because CHCHD2 is a mitochondrial protein involved in oxidative phosphorylation, we examined how METTL5 loss affects cellular metabolism using Seahorse XF metabolic stress tests. Mitochondrial respiration was significantly impaired in *METTL5-*KO NPCs; following sequential injection of oligomycin, FCCP, and rotenone/antimycin A, control NPCs showed the expected dynamic changes in OCR, including a robust increase in maximal respiration after FCCP addition. In contrast, METTL5-KO NPCs had a blunted response to FCCP and markedly reduced maximal and spare respiratory capacity. Quantification of individual time points revealed significant differences at nearly every stage of the assay (**Figure 6A, 6B**; ****p < 0.0001, ***p < 0.001, **p < 0.01, *p < 0.05, ns = not significant, Welch’s t-test). Glycolytic capacity was also diminished in METTL5-KO NPCs, though not as drastically as oxidative respiration (**Supplementary Figure 8**).

**Figure 6:**
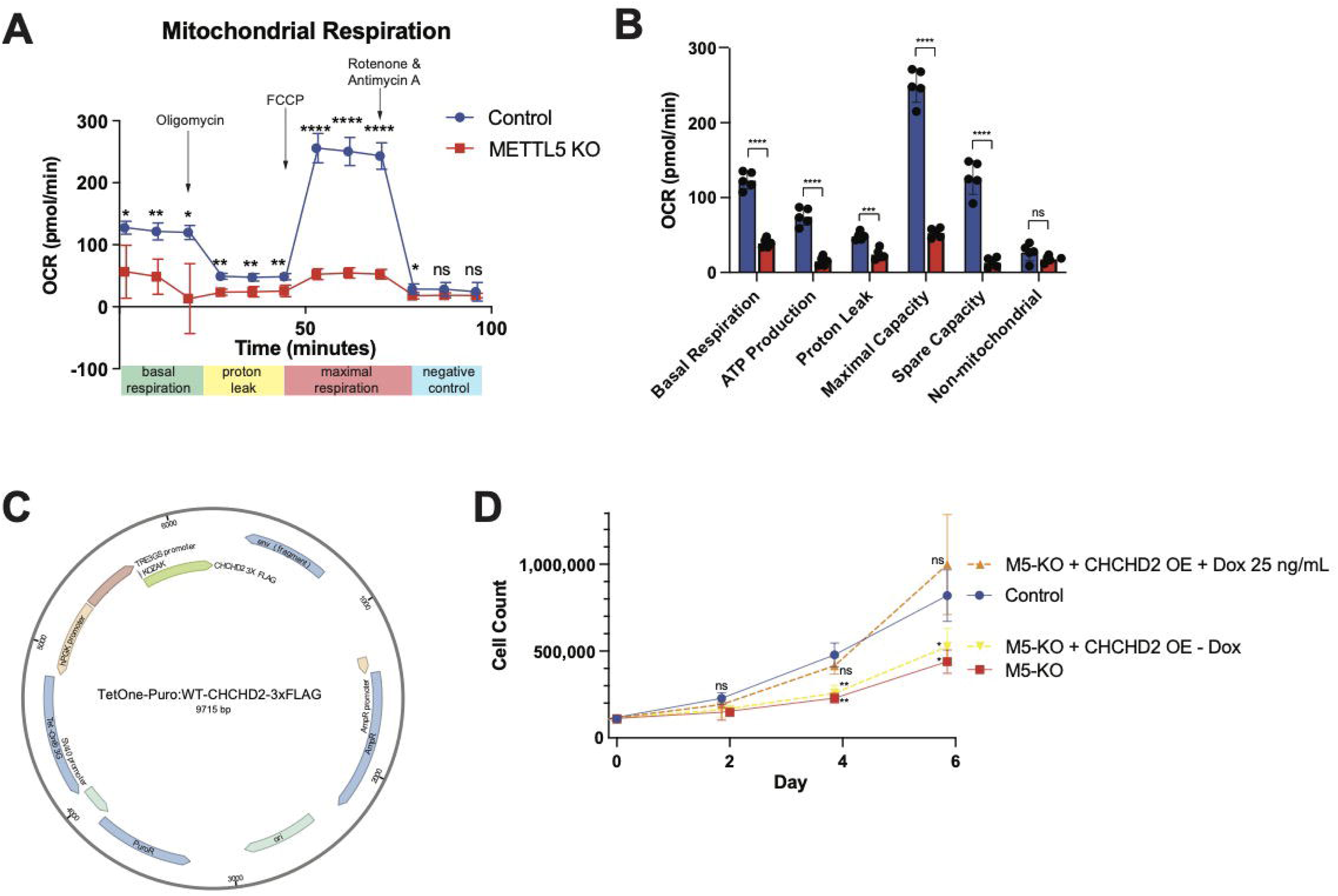
CHCHD2 strongly contributes to impairment of METTL5-KO NPC phenotypes. **A.** Mitochondrial respiration is significantly lower in *METTL5-*KO NPCs at baseline and in response to stress, measured by a Seahorse XF Mito Stress Test. n=5 replicates; *p<0.05, **p<0.01, ****p<0.0001, ns= no significance. **B.** Bar plot quantifications of relative measurements from (**A**). **C.** CHCHD2 overexpression vector with doxycycline (Dox) inducible expression. **D.** Rescue of NPC proliferation rate in *METTL5-KO* CHCHD2 OE NPCs + 25 ng/mL Dox, but not *METTL5-KO* CHCHD2 OE NPCs - Dox. n=3; ns= no significance, *p<0.05, **p<0.01. All significance is relative to control (blue) at each timepoint.

To determine whether the metabolic defects linked to CHCHD2 were significant drivers of the overall phenotype of *METTL5-*KO NPCs, we generated doxycycline-inducible CHCHD2 over expression *METTL5-*KO NPCs (**Figure 6C**). Notably, CHCHD2 overexpression successfully rescues the proliferation rate of *METTL5-*KO NPCs to the level of control NPCs by Day 6 (**Figure 6D**).

## Discussion

In this work, we demonstrate that *METTL5*-KO hiPSC-derived cortical forebrain organoids strongly recapitulate the microcephaly phenotype observed in human patients with METTL5 mutations. Prior studies have reported variable mechanisms for METTL5 functions–some suggesting global regulation of translation, others pointing to transcript-specific effects. These conflicting reports underscore the importance of studying *METTL5* in relevant cellular contexts. Using human cortical organoids, we show that *METTL5* loss significantly impairs neural progenitor cell (NPC) proliferation and neurogenesis, leading to a significant developmental delay. These phenotypes are accompanied by broad translational repression and striking changes in the transcription of neurodevelopmental and metabolic genes.

One of the most notable transcriptional changes in METTL5-KO NPCs is the significant downregulation of CHCHD2, a mitochondrial gene involved in oxidative phosphorylation and cellular stress response. Interestingly, CHCHD2 expression is not altered in *Mettl5*-KO mouse models (*10, 11*), suggesting a possible human-specific role for CHCHD2 in neural development. Supporting this idea, CHCHD2 has recently been implicated in human neurodevelopmental disorders including lissencephaly and autism spectrum disorder (*17*), as well as in neurodegenerative diseases such as Parkinson’s disease and ALS (*19, 31, 32*). Human neural progenitors have high metabolic demands during early development (*33*) and may be particularly sensitive to perturbations in mitochondrial function. Notably, CHCHD2-deficient mouse mitochondria can still maintain oxidative phosphorylation (*34*), highlighting a potential divergence in human mitochondrial vulnerability. In humans, CHCHD2 downregulation disrupts its interaction with its mitochondrial binding partner CHCHD10, leading to impaired oligomerization and downstream dysfunction (*35*). In contrast, mouse CHCHD10 oligomerization appears to be maintained even in the absence of CHCHD2 (*34*), which may help explain species-specific differences in phenotype severity.

Our data show that METTL5-KO NPCs exhibit impaired oxidative metabolism at baseline, with further deficits upon mitochondrial stress. CHCHD2 overexpression in these cells rescues their proliferation defect, suggesting that CHCHD2 loss is a key driver of the observed phenotype in *METTL5-*KO NPCs. The mechanism by which METTL5 regulates CHCHD2 expression remains unclear. Both METTL5 and CHCHD2 have been associated with cellular responses to oxidative stress (*15, 29*), and the neural stem cell niche is known to be hypoxic and under oxidative stress during development (*36, 37*).

While there were no specific system-wide motifs enriched in our translation efficiency data, we did see that the 100 most significantly downregulated subset of transcripts in *METTL5*-KO NPCs were enriched for pyrimidine-rich translation element (PRTE) motifs. These motifs are regulated by mTORC1, with LARP1 binding to 5’TOP and pyrimidine-rich PRTE regions to control ribosome recruitment in an mTOR-dependent manner (*38*). This suggests that METTL5 loss may be intertwined with translational regulation via the mTOR pathway, or that the METTL5-KO translational phenotype is muddied by other forms of translational regulation related to cellular stress. This was also hypothesized in (*13*), which notes that loss of METTL5 in breast cancer cell lines phenocopies MCF7 breast cancer cells treated with mTOR inhibitors. Since mTORC1 activity is closely linked to intracellular ATP levels, impaired oxidative phosphorylation may further suppress mTOR signaling, leading to decreased translation of mTOR-regulated transcripts particularly those containing 5′TOP and PRTE motifs (*38, 39*). This supports a model in which mitochondrial dysfunction and impaired mTOR signaling may partly explain the translational repression observed in METTL5-deficient cells.

Together, these findings highlight a potential mechanism in which METTL5 loss compromises mitochondrial function, leading to broad changes in cellular signaling and gene expression regulation at both transcriptional and translational levels. Future studies are needed to determine whether the transcriptional changes in CHCHD2 and other genes result directly from altered ribosome activity, mitochondrial stress signaling, or a combination of both.

## Methods

### hiPSC Culture

Human iPSCs and hESCs, including KOLF2.1J, H9, and WTC11 lines, were maintained under feeder-free conditions on Matrigel-coated plates. KOLF2.1J and H9 were cultured in StemFlex medium (Thermo Fisher), while WTC11 cells were maintained in mTeSR Plus medium (STEMCELL Technologies). For all lines, culture medium was replenished every other day. Cells were passaged every 4 - 6 days using ReLeSR (STEMCELL Technologies) to selectively detach undifferentiated colonies. Colony morphology was assessed regularly to ensure maintenance of pluripotency, with only compact, defined colonies retained. All lines were karyotyped around 10 passages using the qPCR-based KaryoStat assay (STEMCELL Technologies) to confirm genomic stability.

### hiPSC CRISPR editing

#### Knockout lines

CRISPR-mediated gene knockout was performed in KOLF2.1J, H9, and WTC11 cells using the Synthego Gene Knockout Kit v2, following the manufacturer’s protocol. Chemically modified synthetic single guide RNAs (AGUCGCCUGCAACAAGUGGA, AUUUUUAUCACAGAGUAGAC, GCCGCCCUACCUGCAAUGUG) were designed using Synthego’s online tool to target early exons of the gene of interest and were complexed with recombinant Cas9 protein to form ribonucleoprotein (RNP) complexes. RNPs were delivered into iPSCs via electroporation using the Lonza 4D-Nucleofector system with the P3 Primary Cell Kit. Post-electroporation, cells were plated in StemFlex or mTeSR medium (depending on the cell line) supplemented with 10 µM ROCK inhibitor (Y-27632) to enhance survival. Editing efficiency was assessed 48 - 72 hours post-electroporation by Sanger sequencing and ICE (Inference of CRISPR Edits) analysis. Successfully edited single-cell clones were expanded, genotyped, and validated for indel formation at the target locus.

#### TET-on 3G lines

To reintroduce genes of interest into the METTL5 knockout KOLF2.1J iPSC line, a doxycycline-inducible construct for CHCHD2 was tagged with a C-terminal 3×FLAG epitope and introduced into the METTL5 knockout KOLF2.1J iPSC line using the Tet-On 3G system (Takara Bio). Lentiviral transduction was performed using Takara’s pre-packaged lentiviral components, prepared by reconstituting the 3G packaging mix according to the manufacturer’s instructions.

Knockout iPSCs were transduced with the constructs in the presence of 8 µg/mL polybrene then selected with 0.5µg/mL puromycin for 3 - 5 days. Stable lines were expanded and validated for doxycycline-inducible expression (25Lng/mL) of CHCHD2 by sanger sequencing, qPCR, western blot, and immunofluorescence staining.

#### Proliferation assay

NPCs were grown in a 6 well plate coated with Matrigel (Corning) to approximately 85% confluency before starting the proliferation assay. On Day 0, NPCs were passaged using Accutase (STEMCELL Technologies), spun down at 250g for 5 minutes, and resuspended in Maintenance Medium 1 (MM1) supplemented with 10Lng/mL FGF2, 10Lng/mL EGF, and 10 µM ROCK inhibitor (Y-27632). Cells were counted by hand on a hemacytometer in triplicate to obtain accurate concentrations. 110,000 cells were plated into each well of a 6-well plate coated with Matrigel. Samples were plated in triplicate per timepoint: Days 2, 4, and 6; 9 wells total were seeded for each cell line. 24 hours after seeding, the media was changed to Maintenance Medium 1 (MM1) supplemented with 10Lng/mL FGF2, 10Lng/mL EGF, but without Rock inhibitor. The Day 2 timepoint was collected 48 hours after seeding by removing the media and rinsing each well gently with 1 mL of DPBS. The DPBS was removed and 0.5 mL of Trypsin was added directly into each well, allowed to incubate for 5 minutes at 37°C, and then gently pipetted up and down to generate a single cell suspension. 10 µL of the cells in Trypsin were loaded onto a hemacytometer and counted in duplicates; this was repeated for each of the three replicates per cell line per time point. This collection protocol was repeated for the remaining timepoints. Media was changed on all wells at 24, 48, and 96 hours after seeding.

#### Apoptosis assay

To assess apoptosis, KOLF2.1J neural progenitor cells (NPCs), including both wild-type (WT) and METTL5 knockout (KO) lines, were stained for cleaved caspase-3. For a positive control, cells were treated with 1 μM staurosporine (Sigma-Aldrich) for 4 hours at 37°C to induce apoptosis. Following treatment, cells were fixed in 4% paraformaldehyde (PFA) for 15 minutes at room temperature, then permeabilized with 0.1% Triton X-100 in PBS for 10 minutes. Cells were incubated overnight at 4°C with anti-cleaved caspase-3 antibody (Cell Signaling). The next day, cells were washed and incubated with appropriate Alexa Fluor-conjugated secondary antibodies (1:300, Thermo Fisher) for 2 hours at room temperature. Nuclei were counterstained with DAPI. Imaging was performed using a fluorescence microscope.

#### Seahorse Assay

Mitochondrial function was assessed using the Seahorse XF Mito Stress Test (Agilent Technologies) according to the manufacturer’s instructions. KOLF2.1J wild-type (WT) and METTL5 knockout (KO) neural progenitor cells (NPCs) were seeded into Seahorse XF24 cell culture microplates at a density of [10,000 cells/well] and allowed to adhere overnight in NPC maintenance medium + Ri(10ug/mL). Next day, media was changed to NPC maintenance media. The following day which is the day of the assay, the medium was replaced with Seahorse XF assay medium (Agilent) supplemented with 10 mM glucose, 1 mM pyruvate, and 2 mM glutamine, and cells were incubated at 37°C in a non-CO₂ incubator for 1 hour prior to measurement.

Oxygen consumption rate (OCR) was measured using the Seahorse XFe24 Analyzer following sequential injections of 1 μM oligomycin (ATP synthase inhibitor), 2 μM FCCP (uncoupler), and 0.5 μM rotenone and 0.5 μM antimycin A (complex I and III inhibitors, respectively). Basal respiration, ATP-linked respiration, maximal respiration, spare respiratory capacity, and non-mitochondrial respiration were calculated using Seahorse Wave software. Data were normalized to cell number.

#### Mitochondrial Superoxide assay

Mitochondrial superoxide levels were measured using the MitoBright ROS Deep Red assay (Dojindo Laboratories) according to the manufacturer’s instructions. KOLF2.1J wild-type (WT) and METTL5 knockout (KO) neural progenitor cells (NPCs) were cultured in 35mm glass bottom wells(WPI) until 70% confluency. On the day of the assay, cells were washed once with warm culture medium and incubated with 500 nM MitoBright ROS Deep Red dye diluted in Seahorse XF assay medium (Agilent) supplemented with 10 mM glucose, 1 mM pyruvate, and 2 mM glutamine, for 30 minutes at 37°C in a humidified CO₂ incubator. Nuclei were counterstained with DAPI.

As a positive control for mitochondrial superoxide production, another well of WT and KO cells were treated with 2.5 μM antimycin A (Sigma-Aldrich) for 30 minutes during dye incubation. Following incubation, cells were washed with warm Seahorse XF assay media and imaged live using a fluorescence microscope (excitation/emission: 640/675 nm).

#### Cortical organoid differentiation

**This protocol was adapted from** (***21–24***).

Human pluripotent stem cells (hPSCs) were cultured under feeder-free conditions and dissociated using Accutase (STEMCELL Technologies). For aggregation, 10,000 cells in 100 μL of iPSC maintenance medium supplemented with 10 μM ROCK inhibitor (Y-27632) were seeded into each well of 96-well low-attachment slit-well plates (SBio: MS-9096SZ). Plates were incubated at 37 °C with 5% CO₂; this was designated as Day 0.

On Day 3, a complete media change was performed using Neural Induction Medium, which consisted of GMEM (76.8%; Thermo Fisher Scientific), KnockOut Serum Replacement (20%; Gibco), MEM non-essential amino acids (1X; Thermo Fisher Scientific), sodium pyruvate (1X; Thermo Fisher Scientific), antibiotic-antimycotic (1X; Thermo Fisher Scientific), and 0.18LmM β-mercaptoethanol (Life Technologies). The medium was freshly supplemented with 5 μM SB431542, 3 μM IWR-1-endo, 100 nM LDN-193189 (Bio-Techne/R&D Systems), and 10 μM ROCK inhibitor (Y-27632).

On Day 6, the medium was replaced with Neural Induction Medium containing SB431542, IWR-1-endo, and LDN-193189 but without ROCK inhibitor. From Day 9 to Day 25, complete or partial media changes were performed every other day using Maintenance Medium 1, composed of DMEM/F12 (47.4%) and Neurobasal medium (47.4%) supplemented with B27 minus vitamin A (1X; Thermo Fisher Scientific, 12587010), N2 supplement (1X; Thermo Fisher Scientific), GlutaMAX (1X; Thermo Fisher Scientific), MEM non-essential amino acids (1X; Thermo Fisher Scientific), antibiotic-antimycotic (1X), and 0.1LmM β-mercaptoethanol, freshly supplemented with 10 ng/mL FGF2 and 10 ng/mL EGF.

On Day 18, organoids were transferred to a 37 °C with 5% CO2 set between 57 - 90 rpm, depending on size. From Day 26 to Day 35, medium changes were performed every other day using Maintenance Medium 1 without FGF2 and EGF.

From Day 35 to Day 55, organoids were cultured in Maintenance Medium 2, composed of DMEM/F12 (47.4%) and Neurobasal medium (47.4%) supplemented with B27 with vitamin A (1X; Thermo Fisher Scientific, 17504044), N2 supplement (1X), GlutaMAX (1X), MEM non-essential amino acids (1X), antibiotic-antimycotic (1X), and 0.1LmM β-mercaptoethanol, freshly supplemented with 200LnM ascorbic acid (Innovative Cell Technologies).

From Day 56 to Day 62, medium was replaced with Maintenance Medium 2 supplemented with 200LnM ascorbic acid and 10Lng/mL human LIF (Life Technologies). From Day 63 to Day 70, organoids were cultured in Maintenance Medium 2 with ascorbic acid (200LnM), BDNF (10Lng/mL; PeproTech), and NT-3 (10Lng/mL; Life Technologies). From Day 70 onward, medium changes were performed using Maintenance Medium 2 supplemented only with 200LnM ascorbic acid.

Organoids were harvested on Days 40, 50, 75, and 138. Samples were fixed overnight in 4% paraformaldehyde (PFA) at 4L°C on a shaker set to 60 rpm. The following day, organoids were transferred to 30% sucrose in PBS and incubated overnight at 4L°C with rocking. Organoids were then embedded in Tissue Freezing Medium (TFM; Fisher Scientific) and snap-frozen for downstream analysis.

#### In vitro neurogenesis hiPSC differentiation

This protocol was adapted from (*21–23, 26*).

To induce neural differentiation, human induced pluripotent stem cells (hiPSCs) at 80 - 90% confluency were cultured in Neural Induction Medium (NI medium), composed of GMEM (76.8%; Thermo Fisher Scientific), KnockOut Serum Replacement (20%; Gibco), MEM non-essential amino acids (1X; Thermo Fisher Scientific), sodium pyruvate (1X; Thermo Fisher Scientific), antibiotic-antimycotic (1X; Thermo Fisher Scientific), and 0.18LmM β-mercaptoethanol (Life Technologies). The medium was freshly supplemented with 10 µM SB431542, 100LnM LDN-193189, and 5 µM XAV939 (all from Bio-Techne/R&D Systems).

Medium was changed daily for 7 days. Media volume was adjusted to prevent acidification; additional medium was added if the culture appeared yellow the following day.

By Day 7, healthy neural progenitor cell (NPC) cultures were passaged using Accutase (STEMCELL Technologies) at a 1:3 ratio onto Matrigel-coated (Corning) 6-well plates in Maintenance Medium 1 (MM1) supplemented with 10Lng/mL FGF2, 10Lng/mL EGF, and 10 µM ROCK inhibitor (Y-27632). The following day, medium was replaced with MM1 lacking ROCK inhibitor. NPCs were maintained on Matrigel-coated plates and used for downstream analyses.

For neural differentiation, 800,000 NPCs were seeded into 35Lmm glass-bottom wells coated with poly-L-ornithine (Sigma-Aldrich), laminin-521 (BioLamina), and fibronectin (Corning). MM1 supplemented with ROCK inhibitor was used on the day of seeding, followed by MM1 without ROCK inhibitor the next day. NPCs were maintained in MM1 until Day 9.

On Day 11, the medium was changed to MM1 without supplements. On Day 13, the culture was transitioned to MM2 without supplements. From Day 15 onward, cells were maintained in MM2 supplemented with 500 µM dibutyryl-cAMP (Cell Signaling), 200 µM ascorbic acid (Innovative Cell Technologies), 10Lng/mL BDNF (PeproTech), and 10Lng/mL NT-3 (Life Technologies).

Neurons were collected on Days 10, 20, 30, 40, and 50 for analysis. Cells were fixed in 4% paraformaldehyde (PFA) for 15 minutes at room temperature, washed twice with PBS (Thermo Fisher Scientific), and stored in PBS for downstream staining.

#### hNPC Culture

To induce neural induction, human induced pluripotent stem cells (hiPSCs) at 80 - 90% confluency were cultured in Neural Induction Medium (NI medium), composed of GMEM (76.8%; Thermo Fisher Scientific), KnockOut Serum Replacement (20%; Gibco), MEM non-essential amino acids (1X; Thermo Fisher Scientific), sodium pyruvate (1X; Thermo Fisher Scientific), antibiotic-antimycotic (1X; Thermo Fisher Scientific), and 0.18LmM β-mercaptoethanol (Life Technologies). The medium was freshly supplemented with 10 µM SB431542, 100 nM LDN-193189, and 5 µM XAV939 (all from Bio-Techne/R&D Systems).

Medium was changed daily for 7 days. Media volume was adjusted to prevent acidification; additional medium was added if the culture appeared yellow the following day.

By Day 7, healthy neural progenitor cell (NPC) cultures were passaged using Accutase (STEMCELL Technologies) at a 1:3 ratio onto Matrigel-coated (Corning) 6-well plates in Maintenance Medium 1 (MM1) supplemented with 10 ng/mL FGF2, 10 ng/mL EGF, and 10 µM ROCK inhibitor (Y-27632). The following day, medium was replaced with MM1 lacking ROCK inhibitor. NPCs were maintained on Matrigel-coated plates and used for downstream analyses.

#### qPCR

Total RNA was extracted from cells using the RNeasy Mini Kit (Qiagen) according to the manufacturer’s instructions. RNA concentration and purity were assessed using a NanoDrop spectrophotometer (Thermo Fisher Scientific). For cDNA synthesis, 1 μg of total RNA was reverse transcribed using the SuperScript III Supermix cDNA Synthesis Kit (Invitrogen).

1/10 cDNA samples were used with SYBR® Green Supermix (Thermo Fisher Scientific) according to manufacturer instructions and each reaction was run in technical triplicates. Cycling conditions were as follows: initial denaturation at 95°C for 5 minutes, followed by 39 cycles of 95°C for 10 seconds and 60°C for 30 seconds. A melt curve analysis was performed to confirm primer specificity.

Gene expression was normalized to housekeeping genes such as ACTB, and relative expression levels were calculated using the ΔΔCt method. See table for primers.

#### Table with primers

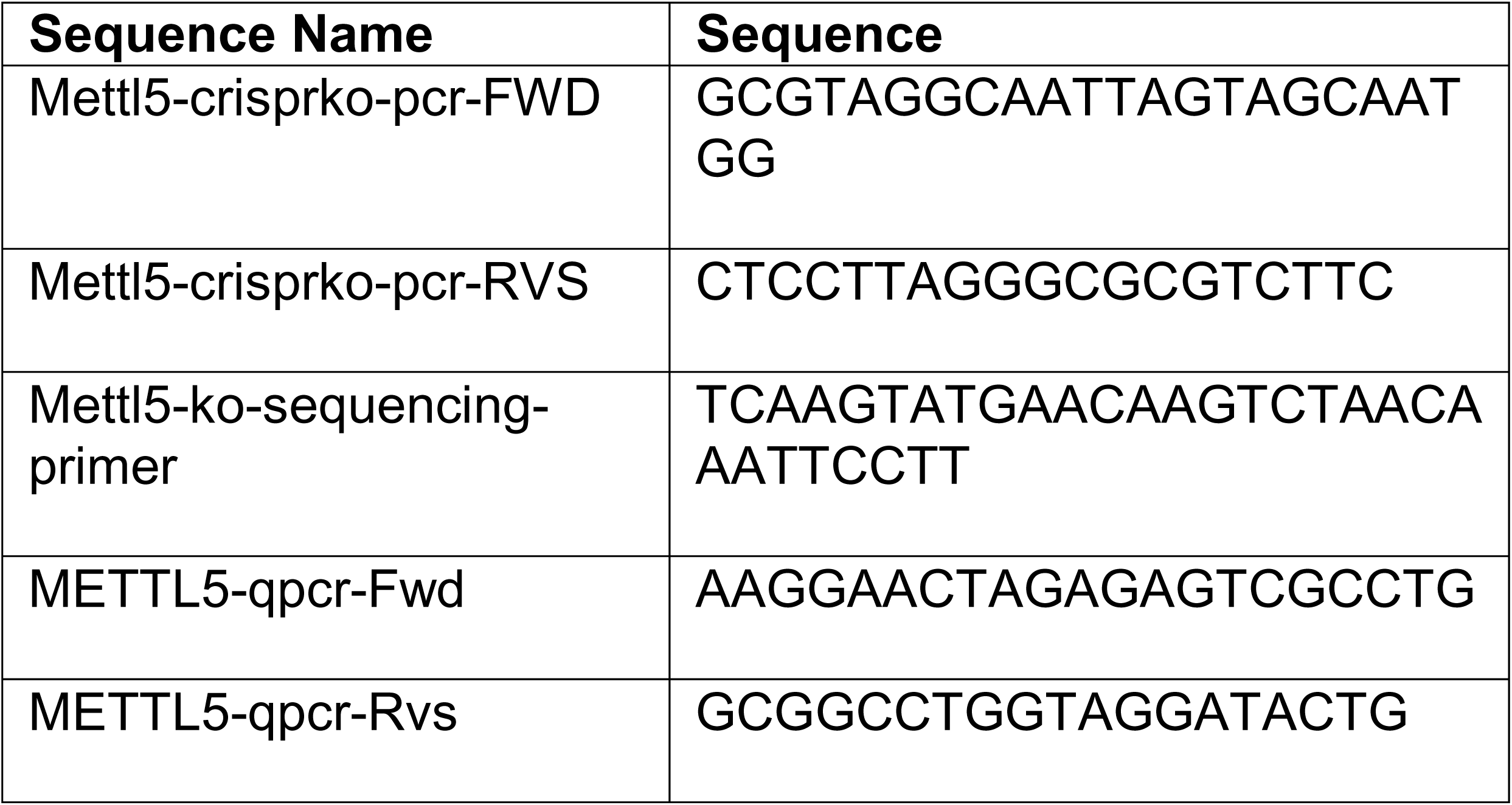
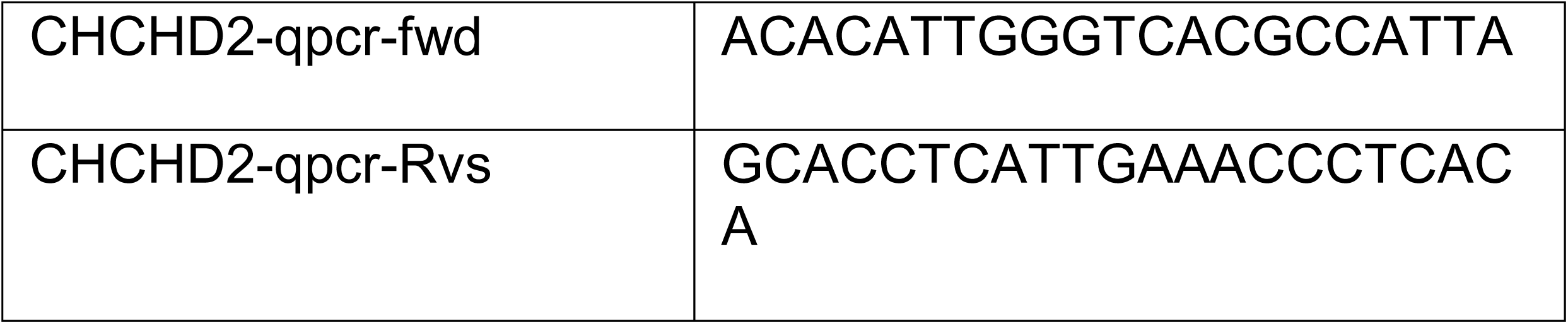

#### Western blot

Cells at 80–90% confluency were lysed in ice-cold RIPA buffer (1X; Thermo Fisher Scientific) supplemented with protease and phosphatase inhibitors (1X; Thermo Fisher Scientific). Prior to lysis, adherent cells were washed twice with ice-cold PBS (Thermo Fisher Scientific). Lysates were incubated on ice for 15-20 minutes and then centrifuged at 12,000 rpm for 20 minutes at 4°C. The resulting supernatants were transferred to pre-chilled 1.5 mL microcentrifuge tubes. Protein concentrations were determined using the Pierce BCA Protein Assay Kit (Thermo Fisher Scientific). Equal amounts of protein (15-20 µg) were mixed with 4X LDS Sample Buffer (Bio-Rad) and 10X NuPAGE Reducing Agent (Thermo Fisher Scientific), then denatured at 70°C for 10 minutes.

Proteins were resolved on 4-12% Bis-Tris gels (Bio-Rad) using 1X NuPAGE MES SDS Running Buffer supplemented with 1X NuPAGE Antioxidant (Life Technologies). Gels were transferred to nitrocellulose membranes using the Power Blotter Select Transfer Stacks and Power Blotter Transfer Buffer (Thermo Fisher Scientific) on a Power Blotter Station. Membranes were blocked in 2% non-fat dry milk and 2% BSA in 1X TBS-T for 1 hour at room temperature with gentle rocking (60 RPM), then incubated overnight at 4°C with primary antibodies diluted in the same blocking buffer. Following three washes in TBS-T, membranes were incubated with HRP-conjugated secondary antibodies (1:5,000 dilution) for 2 hours at room temperature.

Protein bands were detected using the SuperSignal West Dura Extended Duration Substrate (Thermo Fisher Scientific), prepared by mixing equal parts of the enhancer and peroxidase solutions. Chemiluminescence was developed and visualized in the dark.

### Single Cell RNA isolation

#### Organoid dissociation

Wild-type (KOLF2.1J) and METTL5-knockout KOLF2.1J cerebral organoids (three biological replicates per genotype; 3 - 4 organoids pooled per replicate) were dissociated with papain. Organoids were incubated for 15 min at 37 °C in papain (20 U mL-1) activated in EDTA/β-mercaptoethanol/L-cysteine and supplemented with DNase I (0.005 %, w/v) under gentle agitation. Tissue was gently triturated, digested for a further 10 min, then triturated again until a uniform single-cell suspension formed. Digestion was quenched with ovomucoid inhibitor, cells were pelleted (300 x g, 5 min), resuspended in 0.04 % BSA/HBSS, and filtered through a 37 µm strainer. Viability, assessed by trypan blue staining, exceeded 85 % for all samples.

### Single cell RNA seq analysis

#### Library preparation and sequencing

Single cells were processed with the PIPseq-V workflow T10 kit (Fluent BioSciences) according to the manufacturer’s protocol, and libraries were sequenced on an Illumina NovaSeq 6000 (2 × 150 bp).

#### FASTQ processing

Reads were handled with PIPseeker v3.3. Cellular barcodes and intrinsic molecular identifiers (IMIs) were extracted with a Hamming-distance tolerance of 1, written to the read headers, and technical sequences (template-switch oligo, leading T, poly-A) were trimmed.

#### Alignment

Trimmed R2 reads were aligned to the human reference genome GRCh38.p13 (GENCODE v40, Ensembl 106) using STAR v2.7.11a with default PIPseeker settings. Resulting BAM files contain unsorted alignments annotated with barcode (CB) and IMI (UB) tags.

#### Molecule counting

PIPseeker groups reads by cell barcode and gene, collapses identical IMIs that share the same three-base start-site bin, and corrects for amplification bias so that inflated counts are limited to ≤1 %. Multi-mapped molecules are probabilistically assigned to a single gene-barcode pair. The final output is a raw count matrix (matrix.mtx.gz, barcodes.tsv.gz, features.tsv.gz) for downstream analysis.

#### Filtering

Initial quality control was performed to filter out low-quality cells. Cells with fewer than 300 or more than 6,500 detected genes were excluded, as were cells with over 12% mitochondrial gene content, based on thresholds determined from violin plots (Supplementary Figure X). Following QC, each sample was independently normalized using the SCTransform method, regressing out mitochondrial gene content (percent.mt) to correct for technical variation.

Normalized samples were then merged, and metadata was annotated to label wild-type (WT) and knockout (KO) conditions. Principal component analysis (PCA) was performed, and the top 22 principal components were selected based on an elbow plot for downstream analysis. To correct for batch effects across samples, Harmony integration was applied to the selected PCs. Uniform Manifold Approximation and Projection (UMAP) was then used for visualization, and a shared nearest neighbor graph was constructed to perform Louvain clustering.

To determine the optimal clustering resolution, we generated a cluster tree and evaluated cluster stability across resolutions. A resolution of 0.5 was selected for downstream analysis as it provided the best balance between granularity and biological interpretability.

#### Cell type annotation

To annotate cell types, we first optimized the clustering resolution to capture biologically meaningful differences while avoiding over-splitting due to technical variation. We used a cluster tree to evaluate stability across resolutions and selected a resolution of 0.5, which produced clusters corresponding to true cell type markers rather than clusters solely on ribosomal content or cell cycle genes.

Cell type annotations were performed using SCType (*40*), a marker-based annotation tool. We applied SCType with three different gene sets. The annotations were compared across UMAP embeddings to assess consistency in spatial distribution.

To validate and refine these annotations, we used Seurat’s FindAllMarkers function to identify top differentially expressed genes per cluster. We manually examined the expression of canonical markers and referred to the literature for cell type annotations.

#### Differential expression/pseudobulking

To perform differential expression, nuclei were combined into pseudobulk samples per biological replicate using sum aggregation, thus avoiding false positives inherent to nuclei-level pseudoreplication approaches (*41*). The pseudobulk data were then used as input for edgeR glmLRT.

#### GSEA

Cell-type genesets from multiple independent studies and the Broad Institute’s Molecular Signatures Database were used in enrichment analyses (one-sided Fisher’s exact test).

#### rRNA fragment purification

A 40-nt rRNA fragment was purified by performing a previously described protocol (*42*) with slight modifications twice in a row. Briefly, three tubes of total RNA per sample were processed by adding 4 μg of biotinylated DNA probe targeting the A1832 locus of rRNA combined with 33 μg of total RNA in 3.33x hybridization buffer (250mM HEPES pH7, 500mM KCL) in a total volume of 150 uL per tube. The mixtures were incubated for 7 minutes at 90°C and allowed to cool to room temperature over 3.5 hours on the benchtop to promote hybridization. Next, 0.5 μg RNase A (Thermo Scientific) was added to each mixture and incubated for 30 minutes at 37°C to digest unhybridized single-stranded RNA. Mung Bean Nuclease was then added with 10X Mung Bean Nuclease Buffer (NEB) to a concentration of 1X and samples were incubated again for 30 minutes at 37°C to digest any single stranded RNA and DNA. Next, 20 μL of Streptavidin T1 beads (Invitrogen) per sample tube were washed in 1 mL 2.5x IP buffer (375 mM NaCl, 125 mM Tris pH 7.9, 0.25% NP-40) three times using a magnetic rack, then resuspended in 100 μL 2.5x IP buffer. 100 μL of beads were added to each hybridized sample tube and incubated for 1 hr at 4°C on a rotator. The beads were then washed 3 times in 1X IP buffer (150 mM NaCl, 50 mM Tris pH 7.9, 0.1% NP-40), washed twice in nuclease-free water, and then resuspended in 30 uL of nuclease-free water. The resuspended beads were then heated at 70°C for 5 minutes to release the RNA, and immediately placed on ice. The RNA fragment-containing supernatant was collected using a magnetic rack. Supernatants from the three tubes per sample were then combined into one tube per sample and the purification was repeated a second round. The final supernatant collected after the second round of purification was then used for m^6^A quantification.

### m^6^A quantification

#### (CV): purification of 40 nt fragment of 18S rRNA, measure [RNA], perform ELISA according to kit instructions

Double-purified rRNA fragments were used in combination with the EpiQuik™ m^6^A RNA Methylation Quantification Kit (Colorimetric; Epigentek, Farmingdale, NY, USA) to quantify the presence of m^6^A on 18S rRNA A_1832_. Samples were processed according to manufacturer instructions. Briefly, 30 ng of purified rRNA fragment were analyzed alongside a control standard curve (range 0.02 to 1 ng of m^6^A) in a 96 well plate. Signal was detected by reading absorbance at 450 nm with a spectrophotometer CLARIOstar^Plus^ Plate reader (BMG Labtech).

Samples were normalized to a blank well negative control. Sample optical density (OD) was used to calculate the absolute quantity of m^6^A in each sample according to manufacturer instructions:

m^6^A (ng) = (Sample OD-Negative Ctrl OD) / Slope of standard curve

#### Polysome profiling

Control or METTL5 knockout cells were grown on 15cm plates and harvested at 80% confluency and immediately treated with cycloheximide (100 µg/mL) for 3 minutes at 37°C to halt ribosomes on mRNAs. Cells were washed once with ice-cold PBS containing cycloheximide (100 µg/mL) and scraped into a 15cm conical tube. They were spun down at 200xg for 2 minutes then the pellet was lysed in 400ul polysome lysis buffer [20 mM Tris-HCl (pH 7.4), 150 mM NaCl, 5 mM MgCl₂, 1% Triton X-100,, 1 mM DTT, 100 µg/mL cycloheximide, DNAse I 25U/ml), H20], followed by a 20-30 minute incubation on ice. Lysates were clarified by centrifugation at 10,000 X g for 10 minutes at 4°C. RNA was quantified with Nanodrop and 10 - 15ug of RNA was used for the sucrose gradient. Equal amounts of RNA lysate was used for control and knockout.

A 10 - 50% sucrose gradient was prepared in gradient buffer [20 mM Tris-HCl (pH 7.4), 150 mM NaCl, 5 mM MgCl2, 1 mM DTT, and 100 µg/mL cycloheximide] using a BioComp Gradient Master. Equal amounts of clarified lysate (typically 10-15 µg of RNA) were loaded onto the gradient and centrifuged in an SW41 Ti rotor (Beckman Coulter) at 40,000 rpm for 1:45 hours at 4°C.Gradients were fractionated using a BioComp Piston Gradient Fractionator with continuous UV absorbance monitoring at 254 nm to detect ribosome-associated RNA.

Polysome/Monosome A.U.C. Ratio: The area under the curve (AUC) for polysomes was calculated using absorbance values from the 30 - 50% sucrose fractions. The monosome AUC was calculated from the 25 - 29% sucrose fractions. The polysome-to-monosome ratio was determined by dividing the polysome AUC by the monosome AUC. To quantify the relative change, the percent reduction in the ratio was calculated as: ((the ratio of control - ratio of knockout)/ratio of control) X 100.

### Ribosome isolation

#### Footprint isolation

**This protocol was adapted from** (***43–45***)

Control or METTL5 knockout NPCs were grown on 15cm plates (3 plates were combined to make 1 replicate, and control and KO had three replicates each) and harvested at 80% confluency and immediately treated with cycloheximide (100 µg/mL) for 3 minutes at 37°C to halt ribosomes on mRNAs. Cells were washed once with ice-cold PBS containing cycloheximide (100 µg/mL) and lysed in polysome lysis buffer [20 mM Tris-HCl (pH 7.4), 150 mM NaCl, 5 mM MgCl₂, 1% Triton X-100,, 1 mM DTT, 100 µg/mL cycloheximide, DNAse I 25U/ml), H20], followed by a 30 minute incubation on ice. Lysates were clarified by centrifugation at 10,000 X g for 10 minutes at 4°C. 1-2ug of RNA was saved for RNA-seq. 30ug of RNA was incubated with 30U/ml RNAse P1 for 1hr at 37C. The Illustra MicroSpin Columns S-400HR were used for monosome isolation and downstream steps were the same as reported in ()

#### RNA seq

1-2 ug of RNA was purified using the Direct-zol RNA miniprep kit (Zymo) then used in RNA seq library prep kits: NEBNext rRNA Depletion Kit V2 (human/mouse/rat) (NEB), NEBNext Ultra II Directional RNA Library Prep Kit of Illumina (E7760), with NEBNext Multiplex Oligos for Illumina (96 Unique Dual Index Primer Pairs) (E6440S)

### Ribosome sequencing analysis

#### For reads preprocessing and Ribolog

Raw single-end Ribo-seq reads were adapter-trimmed using cutadapt v4.9 with options “--trimmed-only -m 15 -e 0.2 -a TAGACAGATCGGAAGAGCACACGTCTGAACTCCAGTCAC.”

Trimmed reads were then aligned in two steps using bowtie2 v2.5.4. First, reads mapping to a custom rRNA/tRNA index were discarded. The remaining reads were then aligned end-to-end to a GRCh38-based mRNA transcriptome (Ensembl release v96), in which each gene is represented by the isoform containing the longest annotated coding sequences (CDS). The associated total RNA-Seq sample reads were aligned to the same longest CDS reference transcriptome using bowtie2.

#### RNA-seq differential analysis

RNA-seq transcript counts were extracted using the bam2count function in Ribolog. Differential expression analysis was performed using DESeq2 (v.1.42.1) to compare the wild-type and METTL5-knockout samples.

### IGV-like track plot generation

#### Transcriptome track plot generation

To visualize RNA-seq and Ribo-seq coverage profiles, BAM files were first converted to bedGraph files using bedtools genomecov (v2.31.1), normalized by counts per million (CPM). The resulting bedGraph files were used to generate transcriptome-level track plots with the svist4get (v.1.3.1.1) package [https://doi.org/10.1186/s12859-019-2706-8].

#### TE analysis with Ribolog

Ribo-seq and matched RNA-seq reads (BAM files) were imported into Ribolog v0.0.0.9 for differential translation efficiency (TE) analysis. Ribo-seq reads were filtered to retain only in-frame ribosome-protected fragments (RPFs) 40-44 nucleotides long. Codon-level counts derived from these RPFs were corrected for ribosome stalling using the CELP (Consistent Excess of Loess Preds) method implemented in Ribolog. CELP-corrected Ribo-seq and total RNA-seq reads were median-normalized, and lowly expressed transcripts were excluded: specifically, transcripts with fewer than 5 RNA-seq reads in each RNA-seq sample or fewer than 2 RPFs on average across all Ribo-seq samples were removed. TE for each transcript was calculated as the ratio of Ribo-seq to total RNA-seq transcript counts. Differential TE between wild-type and METTL5-knockout samples was modeled using logistic regression and Benjamini–Hochberg correction for multiple testing, as implemented in Ribolog’s logit_seq function.

#### TE analysis with DESeq2

Transcript-level Ribo-seq and RNA-seq counts generated from the Ribolog pipeline were reanalyzed using DESeq2 to assess differential translation efficiency (TE). Ribo-seq counts were obtained from reads filtered for reads with a P-site mapping to the coding sequence (CDS) and a footprint length of 40–44 nucleotides long. RNA-seq counts were extracted using the bam2count function in Ribolog. A two-factor design was used to model differential TE in DESeq2, incorporating sample condition (wild-type or METTL5-knockout) and sequencing assay type (RNA-seq or Ribo-seq), with the following design formula: ∼ assay_type + condition + assay_type:condition.

#### Motif discovery with MEME

To identify regulatory motifs associated with translational changes, genes with significantly altered TE were defined as those overlapping in both Ribolog and DESeq2 analyses (FDR or p-adjusted < 0.05). 5′ untranslated region (5′UTR) sequences were extracted separately for TE-upregulated and TE-downregulated genes. Only 5′UTRs longer than 20 nucleotides were retained. Resulting FASTA files were analyzed using the command-line MEME suite v5.5.5 for de novo motif discovery, with motif widths restricted between 5 and 18 nucleotides.

#### GSEA analysis

Gene set enrichment analysis was performed using the fgsea R package v1.28.0 on differential TE results using the Ribolog method. Genes were ranked by the ratio of the log-fold change to its standard error from the logistic regression model. The fgsea function was used to calculate the enrichment across the MSigDB v2022.1 Hallmark, GO Biological Process, and KEGG pathway gene set collections.

#### Organoid Sectioning and Immunohistochemistry Staining

All organoids were washed in PBS and fixed in 4% PFA at 4°C overnight, then transferred to sterile filtered 30% sucrose in PBS at 4°C for 24 hours. Organoids were then embedded in OCT (TissueTek) and cut to 14-16 µm slices on a Leica CM3050S cryostat. The middle third of organoid sections were prioritized for staining. Brain sections were blocked and permeabilized for 30 minutes at room temperature with blocking solution (5% normal donkey serum, 3% Bovine serum albumin, and 0.1% Triton X-100 in PBS, sterile filtered with 0.44 µm filter. Then, sections were incubated with primary antibodies diluted in the blocking solution at 4°C overnight. Sections were washed 3X in PBST for 10 minutes each at room temperature, then incubated with secondary antibodies diluted in blocking solution for 1 hr at room temperature.

0.1 mg/ml 4,6-diamidino-2-phenylindole (DAPI, Thermo Fisher Scientific) was added to the secondary antibody mixture for visualization of nuclei. Stained sections were mounted with ProLong Gold anti-fade mounting media (Thermo Fisher Scientific) and imaged on a Nikon TE2000 epifluorescent confocal microscope with Zeiss imaging analysis software. Raw images were processed in Fiji ImageJ 1.54p; whole organoid images were stitched using the Stitching Plugin (*46*).

| Antibody | Catalog Number | Concentration |
| --- | --- | --- |
| Ms α Pax6 | Invitrogen MA1-109 | 1:500 |
| Rb α Mettl5 | Atlas HPA-038223 | 1:500 |
| Ms α Nestin | Millipore MAB5326 | 1:500 |
| Ms α Tbr2 | Invitrogen 14-4877-82 | 1:500 |
| Rat α Ctip2 | Abcam 18465 | 1:500 |
| Ms α Tbr1 | Millipore WHOO10716M1 | 1:500 |
| Rb α Caspase 3 | Invitrogen 700182 | 1:500 |

## Acknowledgements

We would like to thank all members of the Oldham lab for thoughtful discussion of results, the Morgan lab for routine support in experiments, and the Fujimori and Goodarzi labs for support with Ribo-sequencing methods and analysis. Sequencing was performed at the UCSF CAT, supported by UCSF PBBR, RRP IMIA, and NIH 1S10OD028511-01 grants. We’d like to acknowledge the UCSF NORC Biomedical Mouse Metabolism & Gnotiobiotics Core supported by the UCSF NORC grant (P30DK098722). This work was also supported by the Program for Breakthrough in Biomedical Research Sandler Foundation Grant (CV) and the UCSF CIRM Scholars Training Program EDUC4-12812 (ET).

---

**Supplementary Figure 1:**
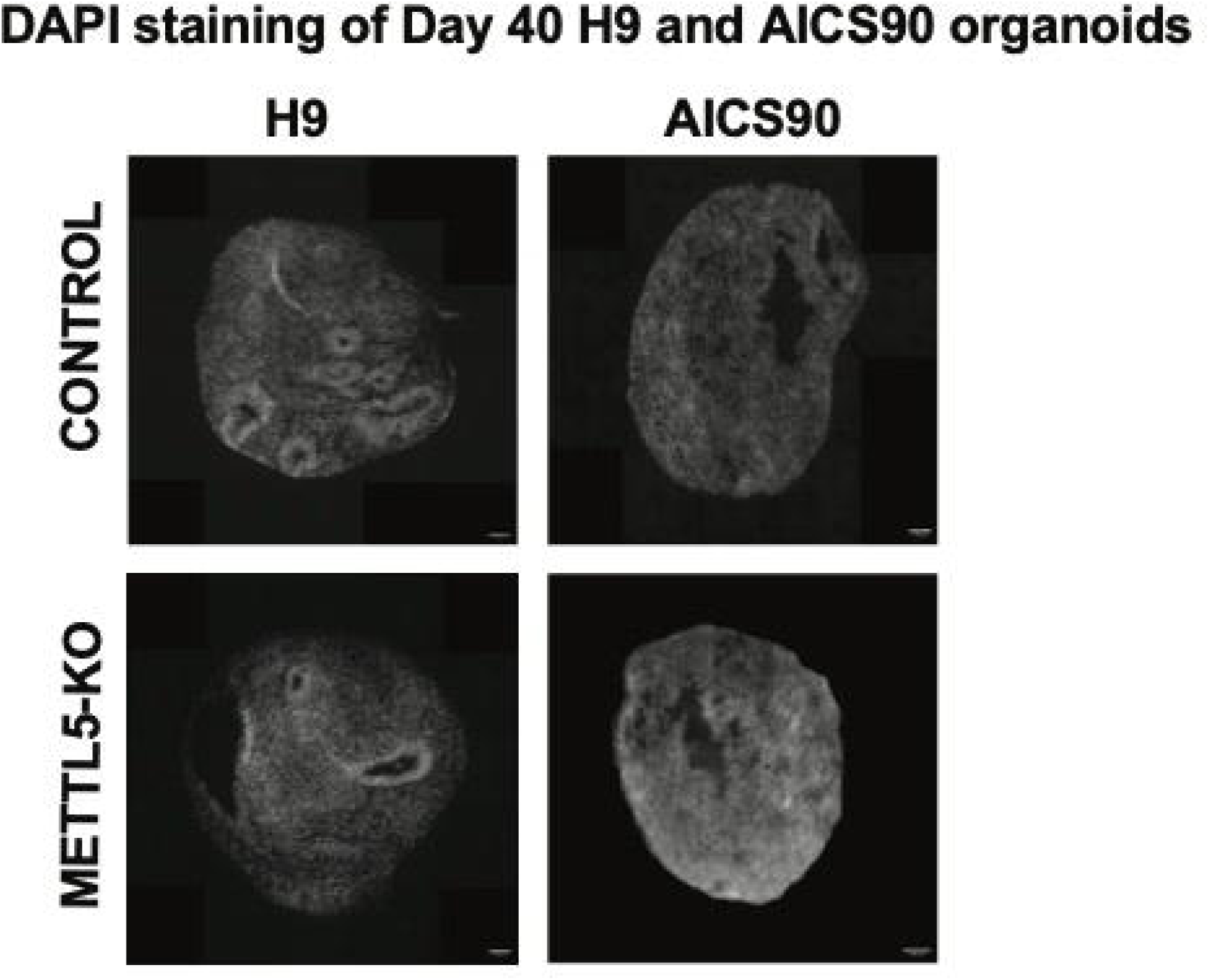
Impairment of neurogenesis in H9 ESC-derived organoids and AICS90 hiPSC-derived organoids at Day 40 in *METTL5*-KO versus controls. DAPI. Scale bar = 100µm

**Supplementary Figure 2:**
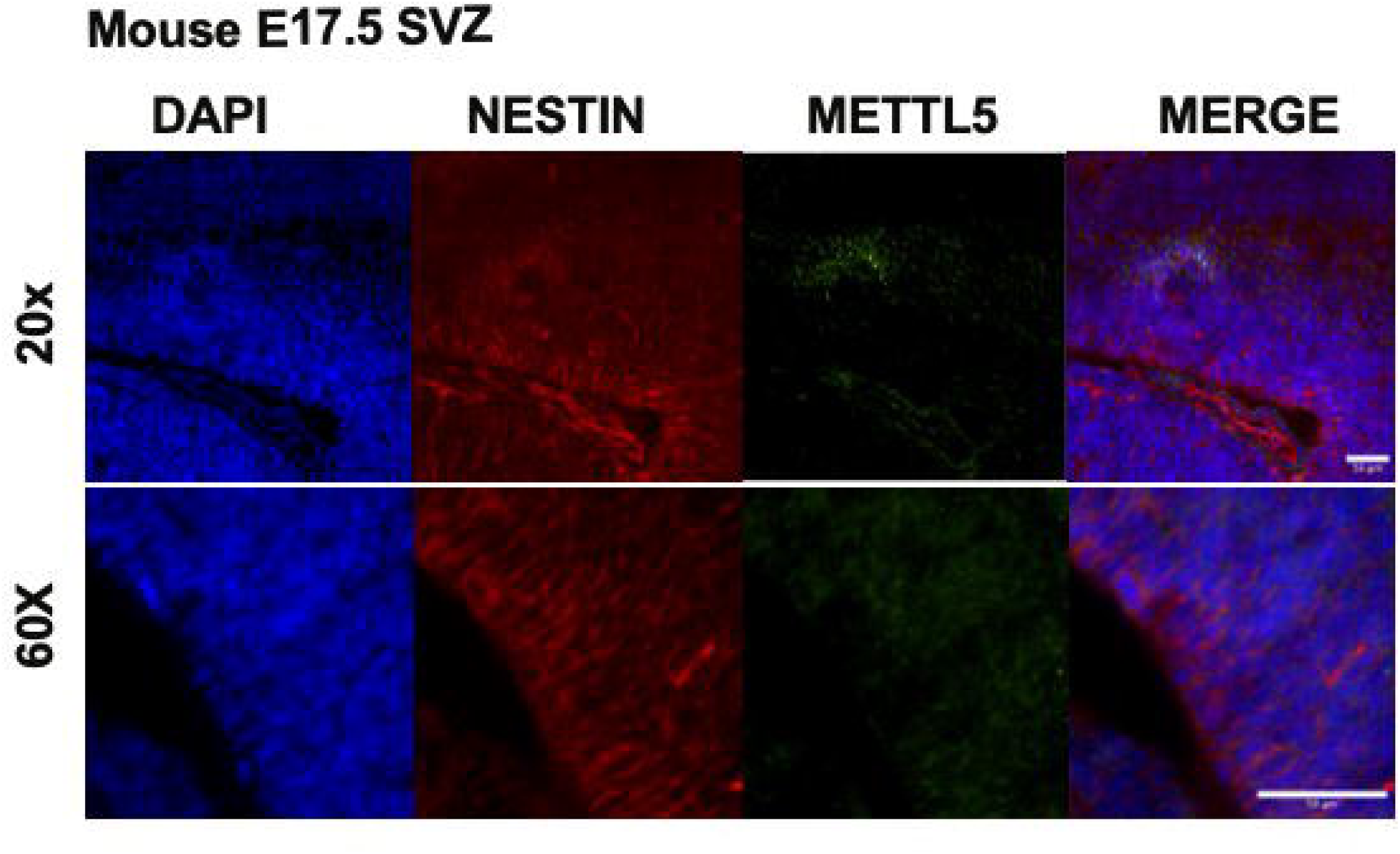
Embryonic day 17.5 mouse ventricle shows Nestin+ radial glial cells in the subventricular zone with minimal METTL5 expression. Scale bars = 50µm

**Supplementary Figure 3:**
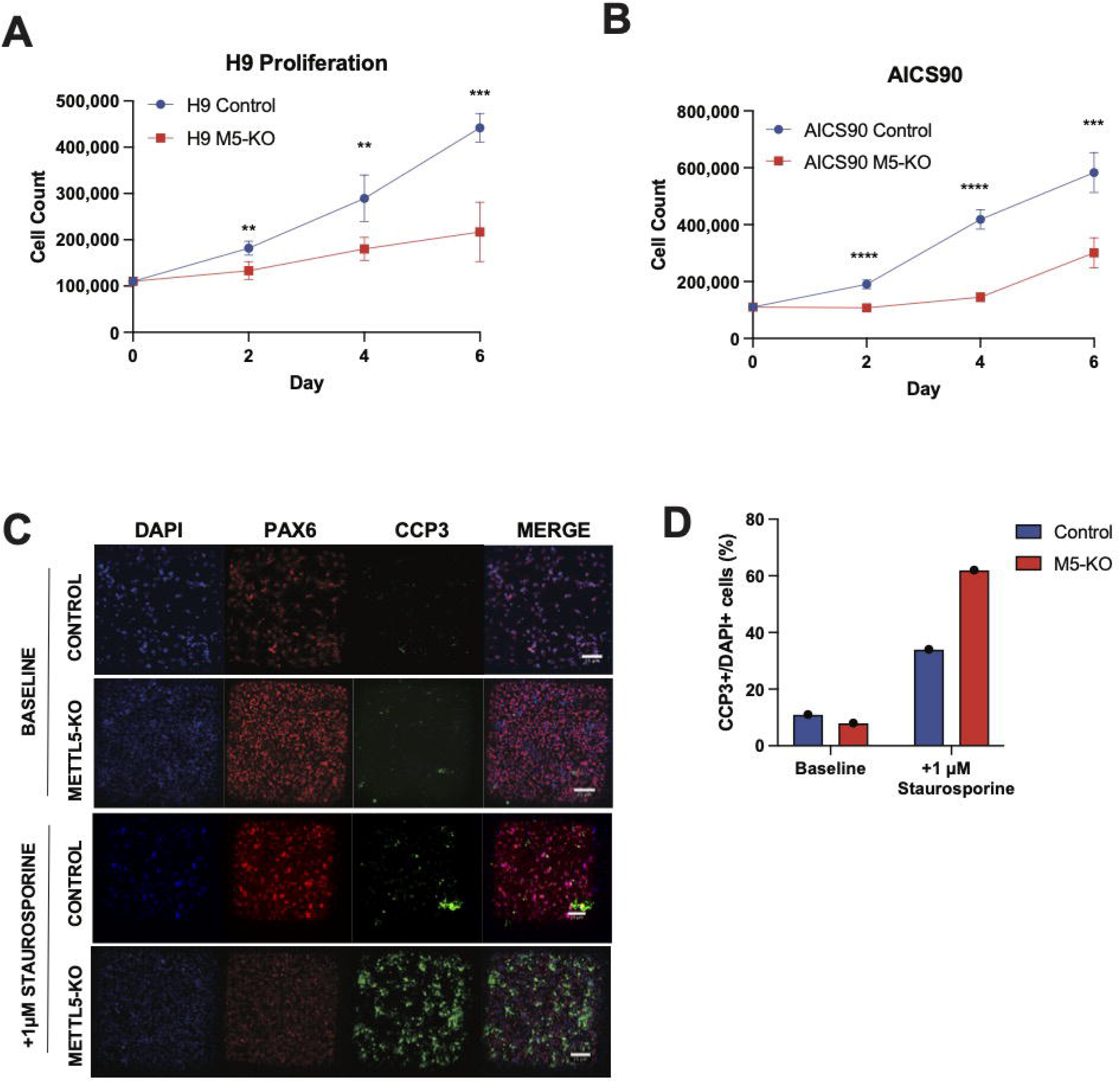
Proliferation is impaired in METTL5-KO NPCs across cell lines. **A.** Recapitulation of impairment in proliferation seen in KOLF2.1J NPCs shown in H9 NPCs and **B.** AICS90 NPCs. n=3; **p<0.01, ***p<0.001, ****p<0.0001. **C.** Changes in proliferation are not due to apoptosis; there is no difference in baseline cleaved caspase 3 (CCP3) levels in control and *METTL5*-KO NPCs, though CCP3 levels are higher in KO NPCs after addition of staurosporine. **D.** Quantification of **(C).**

**Supplementary Figure 4:**
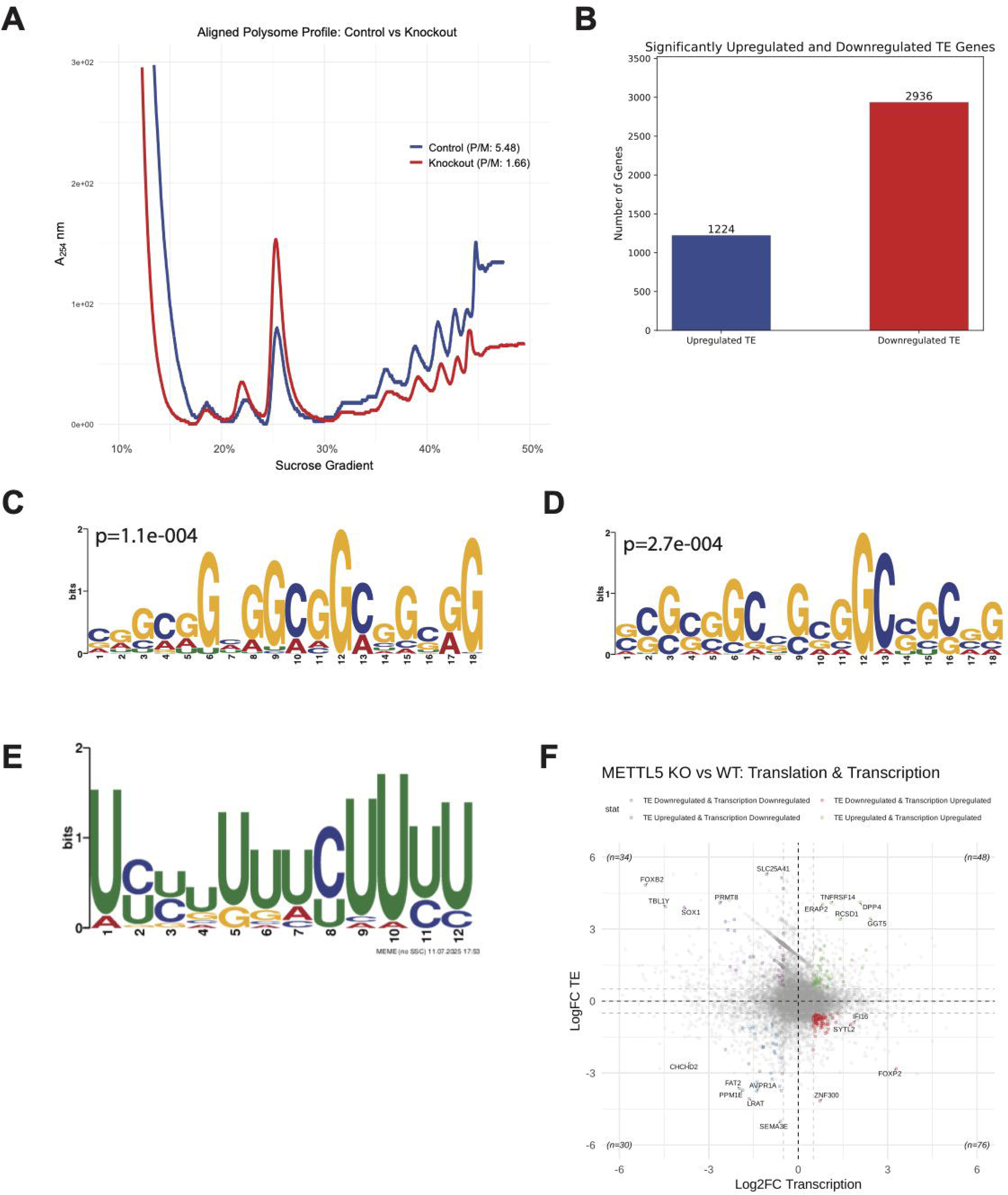
Downregulation of translation in *METTL5-*KO NPCs and iPSCs. **A**. Polysome profiling in KOLF2.1J hiPSCs shows an increase in monosome and decrease in polysomes in *METTL5*-KO hiPSCs compared to control. **B.** Total numbers of significantly up (2936) and down (1224) regulated translation efficiencies in *METTL5*-KO versus control NPCs using n=3 and FDR<0.05. **C.** GC-rich motif enriched in transcripts with significantly higher translation efficiency in *METTL5*-KO/control. **D.** GC-rich motif enriched in transcripts with significantly lower translation efficiency in *METTL5*-KO/control. **E.** Pyrimidine-rich PRTE motif enriched in the top 100 most significantly (FDR) downregulated genes in *METTL5-KO*/Control translation efficiency. p=4.1e-002 **F.** Plot of Log Fold_2_ Change in translation efficiency (y-axis) measured by RiboLog versus Log Fold_2_ Change in transcription (DESeq) show no clear correlation between transcriptional and translational differential regulation.

**Supplementary Figure 5:**
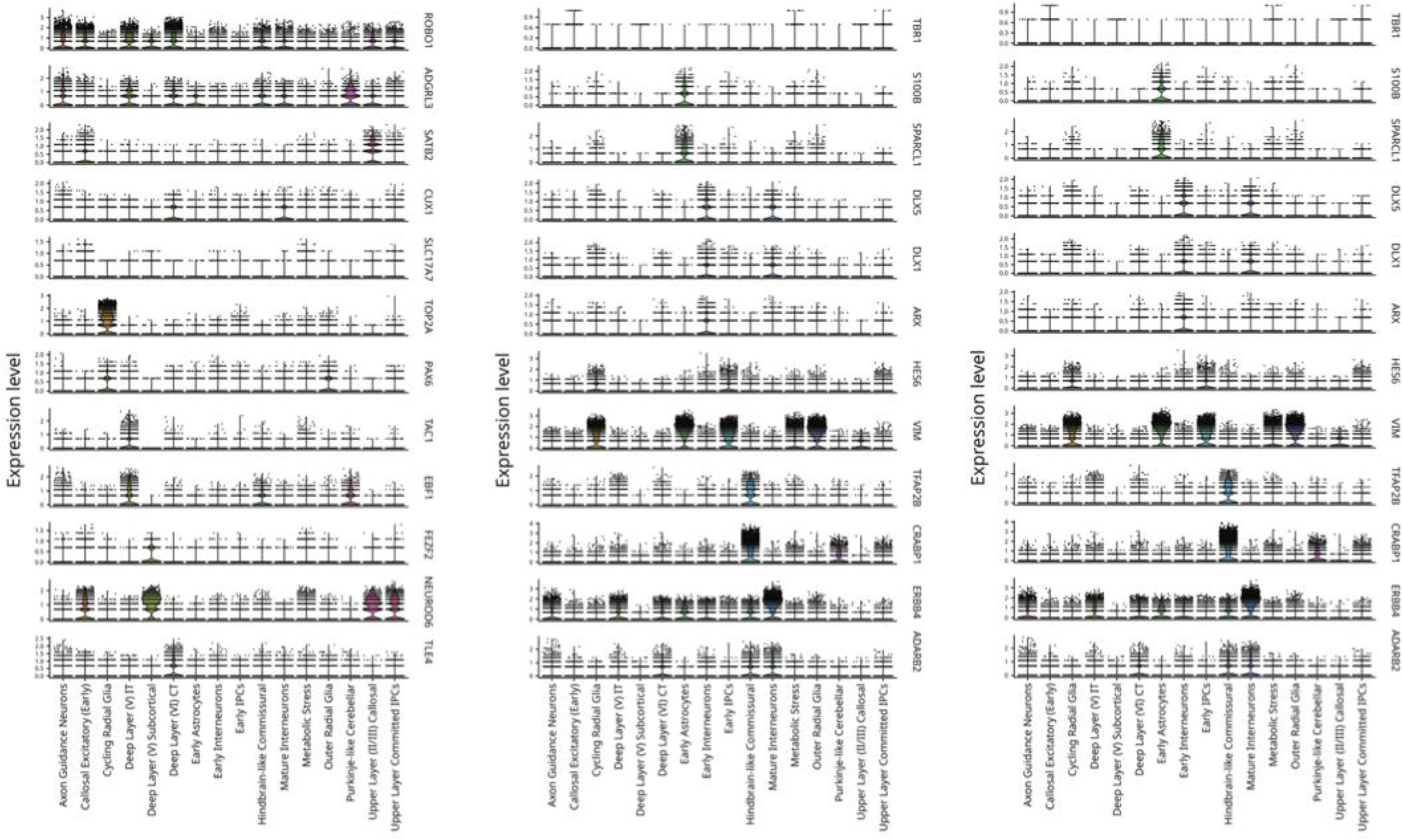
Feature plots of representative markers for each cell type identified in scRNA-seq of Day 75 organoids.

**Supplementary Figure 6:**
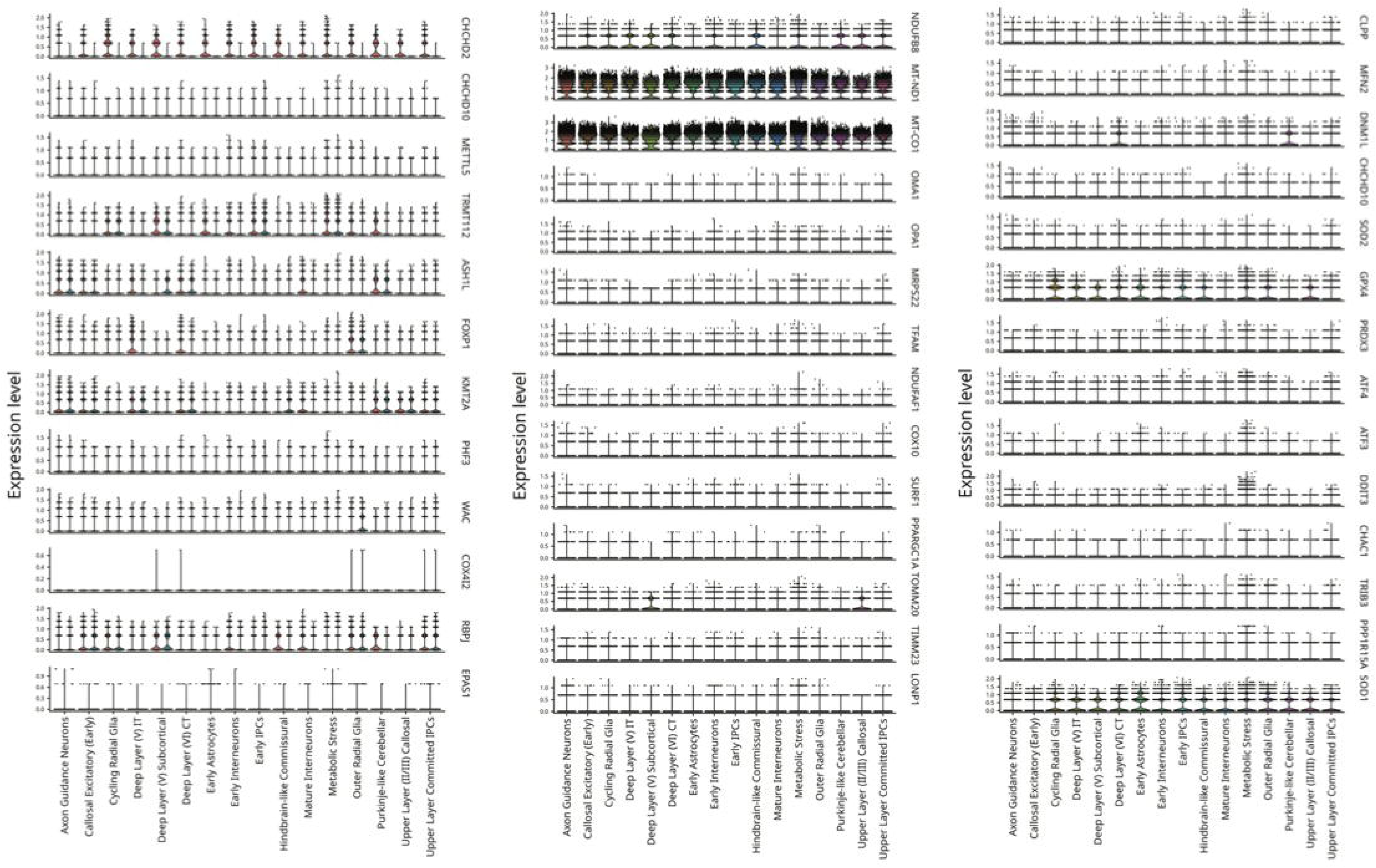
Feature plots of markers of interest in each cell type identified.

**Supplementary Figure 7:**
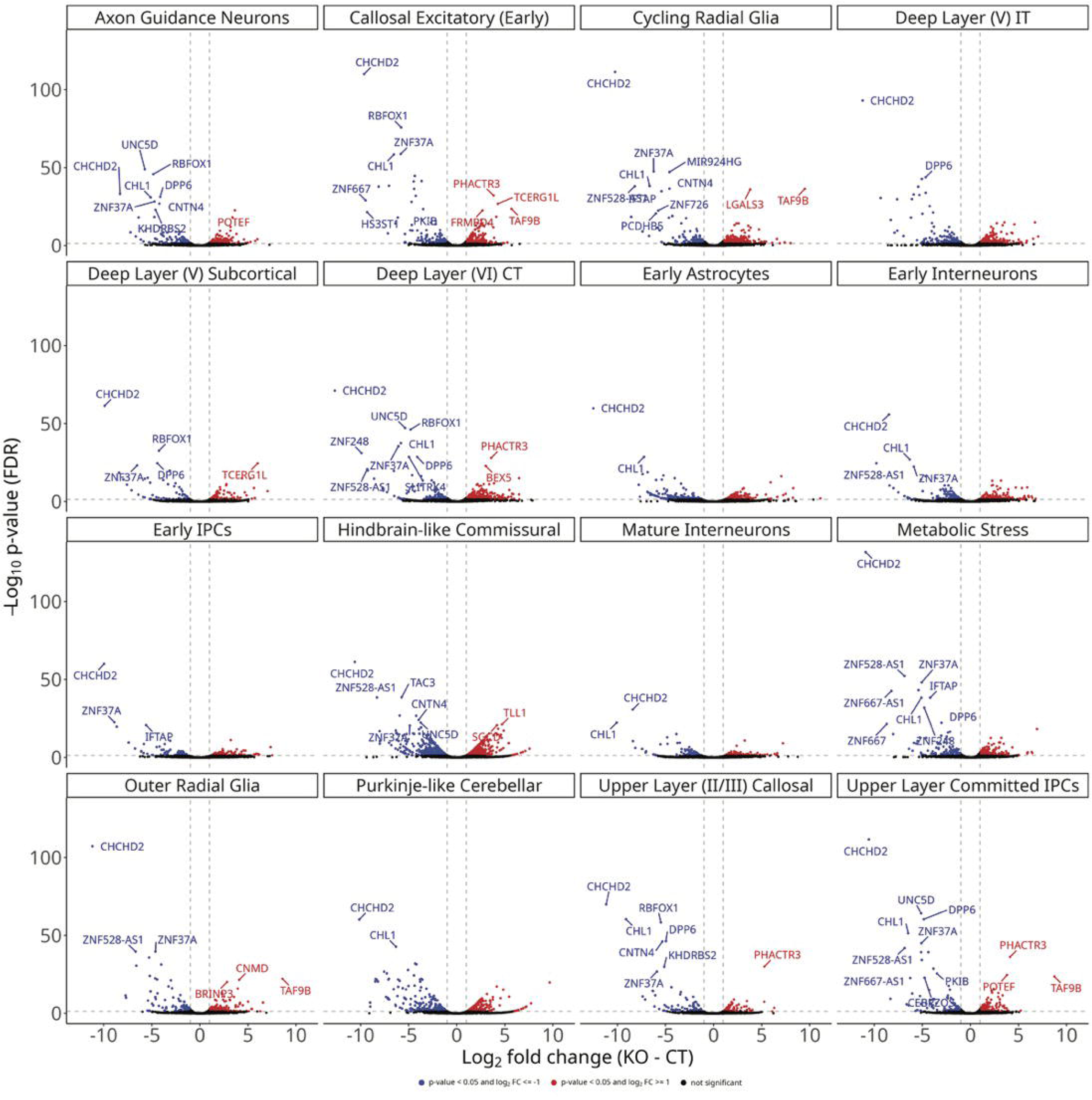
Volcano plots of differentially expressed genes between control and *METTL5-*KO Day 75 organoids in individual cell clusters from Figure 5a.

**Supplementary Figure 8:**
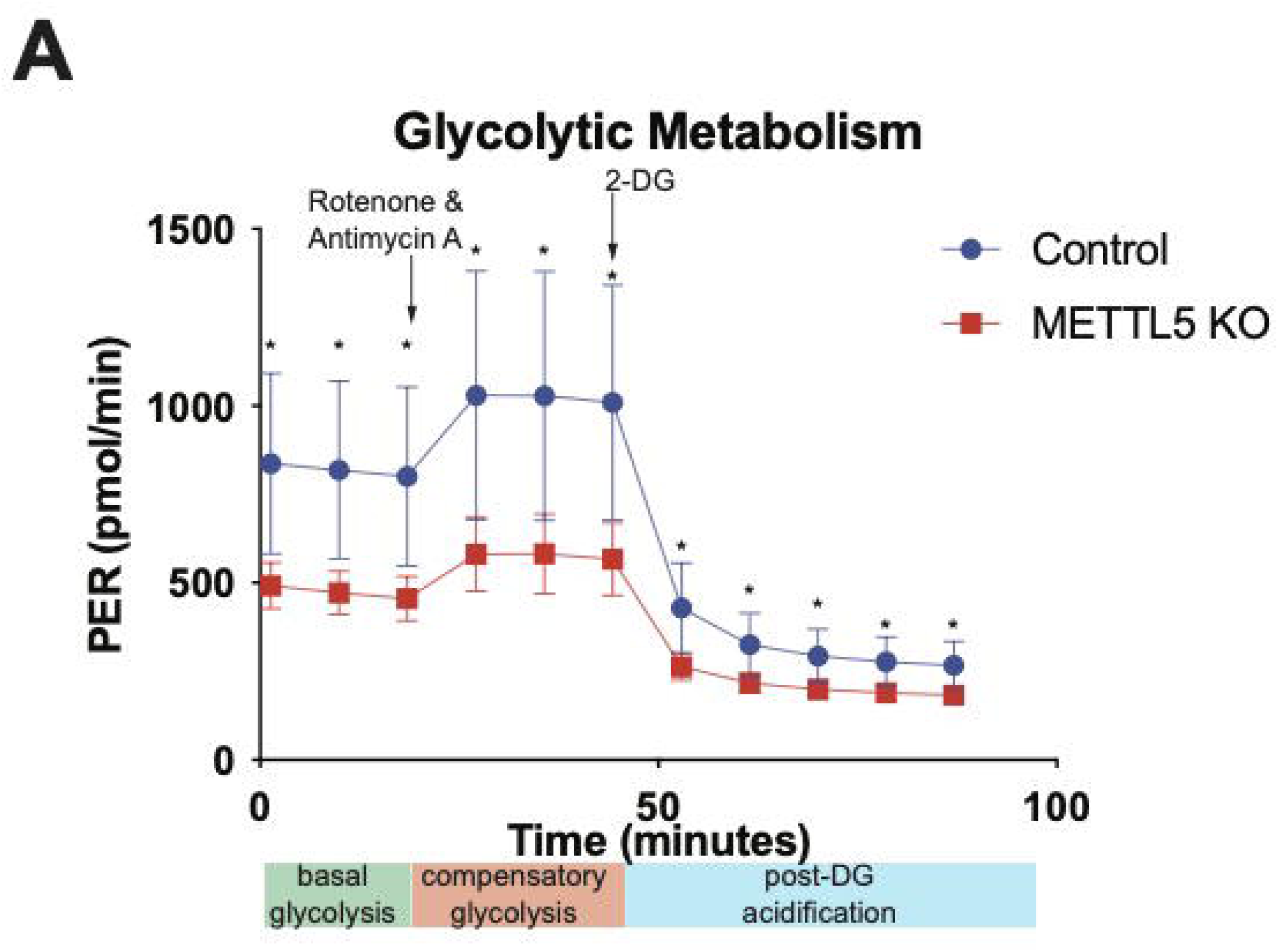
*METTL5-KO* NPCs show minor impairments in glycolytic metabolism. n=5; *p<0.05

## Notes

### Competing Interest Statement

The authors have declared no competing interest.

## Works Cited

1. N. van Tran et al., The human 18S rRNA m6A methyltransferase METTL5 is stabilized by TRMT112. Nucleic Acids Res 47, 7719–7733 (2019).

2. E. M. Turkalj, C. Vissers, The emerging importance of METTL5-mediated ribosomal RNA methylation. Exp Mol Med 54, 1617–1625 (2022).

3. E. M. Richard et al., Bi-allelic Variants in METTL5 Cause Autosomal-Recessive Intellectual Disability and Microcephaly. Am J Hum Genet 105, 869–878 (2019).

4. D. Torun, M. Arslan, B. Cavdarli, H. Akar, D. S. Cram, Three Afghani siblings with a novel homozygous variant and further delineation of the clinical features of METTL5 related intellectual disability syndrome. Turk J Pediatr 64, 956–963 (2022).

5. F. Shakarami, Z. Nouri, H. Khanahmad, M. Ghazavi, M. Amin Tabatabaiefar, A novel METTL5 variant disrupting a donor splice site leads to primary microcephaly-related intellectual disability in an Iranian family: clinical features and literature review. J Genet 102, (2023).

6. X. Zhou et al., Microcephaly-related global developmental delay caused by a pathogenic METTL5 splicing mutation in a Chinese family. J Hum Genet, (2025).

7. M. DeSilva et al., Congenital microcephaly: Case definition & guidelines for data collection, analysis, and presentation of safety data after maternal immunisation. Vaccine 35, 6472–6482 (2017).

8. V. V. Ignatova et al., The rRNA m(6)A methyltransferase METTL5 is involved in pluripotency and developmental programs. Genes Dev 34, 715–729 (2020).

9. K. Lei et al., METTL5 regulates cranial suture fusion via Wnt signaling. Fundam Res 3, 369–376 (2023).

10. C. Sepich-Poore et al., The METTL5-TRMT112 N(6)-methyladenosine methyltransferase complex regulates mRNA translation via 18S rRNA methylation. J Biol Chem 298, 101590 (2022).

11. L. Wang et al., Mettl5 mediated 18S rRNA N6-methyladenosine (m(6)A) modification controls stem cell fate determination and neural function. Genes Dis 9, 268–274 (2022).

12. J. Leismann et al., The 18S ribosomal RNA m(6) A methyltransferase Mettl5 is required for normal walking behavior in Drosophila. EMBO Rep 21, e49443 (2020).

13. B. Rong et al., Ribosome 18S m(6)A Methyltransferase METTL5 Promotes Translation Initiation and Breast Cancer Cell Growth. Cell Rep 33, 108544 (2020).

14. M. Xing et al., The 18S rRNA m(6) A methyltransferase METTL5 promotes mouse embryonic stem cell differentiation. EMBO Rep 21, e49863 (2020).

15. Q. L. Hao Chen, Dan Yu, Kundhavai Natchiar, Chen Zhou, Chih-hung Hsu, Pang-Hung Hsu, Xing Zhang, Bruno Klaholz, Richard I. Gregory, Xiaodong Cheng, Yang Shi, METTL5, an 18S rRNA-specific m6A methyltransferase, modulates expression of stress response genes. bioRxiv, (2020).

16. K. Shimojima et al., CHCHD2 is down-regulated in neuronal cells differentiated from iPS cells derived from patients with lissencephaly. Genomics 106, 196–203 (2015).

17. C. Li et al., Single-cell brain organoid screening identifies developmental defects in autism. Nature 621, 373–380 (2023).

18. G. Gao, Y. Shi, H. X. Deng, D. Krainc, Dysregulation of mitochondrial alpha-ketoglutarate dehydrogenase leads to elevated lipid peroxidation in CHCHD2-linked Parkinson’s disease models. Nat Commun 16, 1982 (2025).

19. P. Lisowski et al., Mutant huntingtin impairs neurodevelopment in human brain organoids through CHCHD2-mediated neurometabolic failure. Nat Commun 15, 7027 (2024).

20. Y. Huang et al., CHCHD2 rescues the mitochondrial dysfunction in iPSC-derived neurons from patient with Mohr-Tranebjaerg syndrome. Cell Death Dis 16, 173 (2025).

21. T. Kadoshima et al., Self-organization of axial polarity, inside-out layer pattern, and species-specific progenitor dynamics in human ES cell-derived neocortex. Proc Natl Acad Sci U S A 110, 20284–20289 (2013).

22. M. Bershteyn et al., Human iPSC-Derived Cerebral Organoids Model Cellular Features of Lissencephaly and Reveal Prolonged Mitosis of Outer Radial Glia. Cell Stem Cell 20, 435–449 e434 (2017).

23. R. S. Ziffra et al., Single-cell epigenomics reveals mechanisms of human cortical development. Nature 598, 205–213 (2021).

24. T. Bertucci et al., Improved Protocol for Reproducible Human Cortical Organoids Reveals Early Alterations in Metabolism with MAPT Mutations. bioRxiv, (2023).

25. C. M. M. Karch, “Differentiation of NPC into cortical neurons,” (2019).

26. Y. Qi et al., Combined small-molecule inhibition accelerates the derivation of functional cortical neurons from human pluripotent stem cells. Nat Biotechnol 35, 154–163 (2017).

27. J. D. Blair, D. Hockemeyer, J. A. Doudna, H. S. Bateup, S. N. Floor, Widespread Translational Remodeling during Human Neuronal Differentiation. Cell Rep 21, 2005–2016 (2017).

28. A. Navickas et al., An mRNA processing pathway suppresses metastasis by governing translational control from the nucleus. Nat Cell Biol 25, 892–903 (2023).

29. Y. Ruan et al., CHCHD2 and CHCHD10 regulate mitochondrial dynamics and integrated stress response. Cell Death Dis 13, 156 (2022).

30. L. Zhu et al., The mitochondrial protein CHCHD2 primes the differentiation potential of human induced pluripotent stem cells to neuroectodermal lineages. J Cell Biol 215, 187–202 (2016).

31. X. Liu et al., CHCHD2 up-regulation in Huntington disease mediates a compensatory protective response against oxidative stress. Cell Death Dis 15, 126 (2024).

32. C. H. Shi et al., CHCHD2 gene mutations in familial and sporadic Parkinson’s disease. Neurobiol Aging 38, 217 e219–217 e213 (2016).

33. X. Zheng et al., Metabolic reprogramming during neuronal differentiation from aerobic glycolysis to neuronal oxidative phosphorylation. Elife 5, (2016).

34. M. K. Nguyen et al., Mouse midbrain dopaminergic neurons survive loss of the PD-associated mitochondrial protein CHCHD2. Hum Mol Genet 31, 1500–1518 (2022).

35. X. Huang et al., CHCHD2 accumulates in distressed mitochondria and facilitates oligomerization of CHCHD10. Hum Mol Genet 28, 349 (2019).

36. A. Mohyeldin, T. Garzon-Muvdi, A. Quinones-Hinojosa, Oxygen in stem cell biology: a critical component of the stem cell niche. Cell Stem Cell 7, 150–161 (2010).

37. J. A. Ortega, C. L. Sirois, F. Memi, N. Glidden, N. Zecevic, Oxygen Levels Regulate the Development of Human Cortical Radial Glia Cells. Cereb Cortex 27, 3736–3751 (2017).

38. T. Hochstoeger, P. Papasaikas, E. Piskadlo, J. A. Chao, Distinct roles of LARP1 and 4EBP1/2 in regulating translation and stability of 5’TOP mRNAs. Sci Adv 10, eadi7830 (2024).

39. M. Morita et al., mTORC1 controls mitochondrial activity and biogenesis through 4E-BP-dependent translational regulation. Cell Metab 18, 698–711 (2013).

40. K. Lanevskij, R. Didziapetris, A. Sazonovas, Physicochemical QSAR analysis of hERG inhibition revisited: towards a quantitative potency prediction. J Comput Aided Mol Des 36, 837–849 (2022).

41. A. E. Murphy, N. G. Skene, A balanced measure shows superior performance of pseudobulk methods in single-cell RNA-sequencing analysis. Nat Commun 13, 7851 (2022).

42. H. Ma et al., N(6-)Methyladenosine methyltransferase ZCCHC4 mediates ribosomal RNA methylation. Nat Chem Biol 15, 88–94 (2019).

43. B. Bibel et al., Context-specific inhibition of mitochondrial ribosomes by phenicol and oxazolidinone antibiotics. Nucleic Acids Res 53, (2025).

44. Z. Jowhar et al., A ubiquitous GC content signature underlies multimodal mRNA regulation by DDX3X. Mol Syst Biol 20, 276–290 (2024).

45. N. J. McGlincy, N. T. Ingolia, Transcriptome-wide measurement of translation by ribosome profiling. Methods 126, 112–129 (2017).

46. S. Preibisch, S. Saalfeld, P. Tomancak, Globally optimal stitching of tiled 3D microscopic image acquisitions. Bioinformatics 25, 1463–1465 (2009).

